# Cytoarchitectonic gradients of laminar degeneration in behavioral variant frontotemporal dementia

**DOI:** 10.1101/2024.04.05.588259

**Authors:** Daniel T. Ohm, Sharon X. Xie, Noah Capp, Sanaz Arezoumandan, Katheryn A.Q. Cousins, Katya Rascovsky, David A. Wolk, Vivianna M. Van Deerlin, Edward B. Lee, Corey T. McMillan, David J. Irwin

**Author notes:** Please send correspondence to: Daniel T. Ohm, PhD, Penn Digital Neuropathology Lab, Frontotemporal Degeneration Center University of Pennsylvania Perelman School of Medicine.

## Abstract

Behavioral variant frontotemporal dementia (bvFTD) is a clinical syndrome primarily caused by either tau (bvFTD-tau) or TDP-43 (bvFTD-TDP) proteinopathies. We previously found lower cortical layers and dorsolateral regions accumulate greater tau than TDP-43 pathology; however, patterns of laminar neurodegeneration across diverse cytoarchitecture in bvFTD is understudied. We hypothesized that bvFTD-tau and bvFTD-TDP have distinct laminar distributions of pyramidal neurodegeneration along cortical gradients, a topologic order of cytoarchitectonic subregions based on increasing pyramidal density and laminar differentiation. Here, we tested this hypothesis in a frontal cortical gradient consisting of five cytoarchitectonic types (i.e., periallocortex, agranular mesocortex, dysgranular mesocortex, eulaminate-I isocortex, eulaminate-II isocortex) spanning anterior cingulate, paracingulate, orbitofrontal, and mid-frontal gyri in bvFTD-tau (n=27), bvFTD-TDP (n=47), and healthy controls (HC; n=32). We immunostained all tissue for total neurons (NeuN; neuronal-nuclear protein) and pyramidal neurons (SMI32; non-phosphorylated neurofilament) and digitally quantified NeuN-immunoreactivity (ir) and SMI32-ir in supragranular II-III, infragranular V-VI, and all I-VI layers in each cytoarchitectonic type. We used linear mixed-effects models adjusted for demographic and biologic variables to compare SMI32-ir between groups and examine relationships with the cortical gradient, long-range pathways, and clinical symptoms. We found regional and laminar distributions of SMI32-ir expected for HC, validating our measures within the cortical gradient framework. While SMI32-ir loss was not related to the cortical gradient in bvFTD-TDP, SMI32-ir progressively decreased along the cortical gradient of bvFTD-tau and included greater SMI32-ir loss in supragranular eulaminate-II isocortex in bvFTD-tau vs bvFTD-TDP (*p*=0.039). In a structural model for long-range laminar connectivity between infragranular mesocortex and supragranular isocortex, we found a larger laminar ratio of mesocortex-to-isocortex SMI32-ir in bvFTD-tau vs bvFTD-TDP (*p*=0.019), suggesting select long-projecting pathways may contribute to isocortical-predominant degeneration in bvFTD-tau. In cytoarchitectonic types with the highest NeuN-ir, we found lower SMI32-ir in bvFTD-tau vs bvFTD-TDP (*p*=0.047), suggesting pyramidal neurodegeneration may occur earlier in bvFTD-tau. Lastly, we found that reduced SMI32-ir related to behavioral severity and frontal-mediated letter fluency, not temporal-mediated confrontation naming, demonstrating the clinical relevance and specificity of frontal pyramidal neurodegeneration to bvFTD-related symptoms. Our data suggest loss of neurofilament-rich pyramidal neurons is a clinically relevant feature of bvFTD that selectively worsens along a frontal cortical gradient in bvFTD-tau, not bvFTD-TDP. Therefore, tau-mediated degeneration may preferentially involve pyramidal-rich layers that connect more distant cytoarchitectonic types. Moreover, the hierarchical arrangement of cytoarchitecture along cortical gradients may be an important neuroanatomical framework for identifying which types of cells and pathways are differentially involved between proteinopathies.

## INTRODUCTION

The behavioral variant of frontotemporal dementia (bvFTD) is a clinical syndrome characterized by progressive and heterogeneous changes in social behavior, cognition, and personality.^1^ Cognitive and behavioral symptoms are related to patterns of frontotemporal-predominant cortical atrophy and selective vulnerability of salience, semantic, and executive networks in bvFTD.^2–7^ A range of frontotemporal lobar degeneration (FTLD) proteinopathies can cause bvFTD,^8^ including a similar incidence of transactive response DNA-binding protein of 43kDa (TDP-43) pathology (FTLD-TDP) and FTLD-type tau pathology (FTLD-tau) among bvFTD patients.^8–13^ This clinicopathologic heterogeneity, in addition to limited biomarkers, currently hinder reliable antemortem diagnosis of the causative FTLD-type pathology in bvFTD.^14^ Postmortem neuroanatomical changes associated with proteinopathies in bvFTD remain poorly understood. However, there is growing evidence that there are different neuroanatomical loci of peak pathologic aggregation and subsequent neurodegeneration in bvFTD, implicating different cellular pathways in the bvFTD spectrum. For example, distinct distributions of cortical atrophy indicate there are anatomical subtypes of bvFTD that interestingly share early behavioral features but differ in other cognitive domains such as executive functioning.^15–17^ MRI-based anatomical variants of bvFTD have not yet directly linked to specific FTLD pathologic/genetic subtypes, but autopsy-confirmed comparative studies have the unique opportunity to address this gap in understanding.

Indeed, postmortem examinations of neuropathology and neurodegeneration can be conducted in cytoarchitectonic subregions and cellular layers not yet resolvable by MRI. At the level of regional cortices, we reported that bvFTD patients accumulate greater TDP-43 pathology in orbitofrontal cortices and greater tau pathology in more dorsal areas including mid-frontal and anterior cingulate cortices.^18–20^ At the level of cortical layers, we recently found preferential accumulation of TDP-43 pathology in upper layers in comparison to preferential accumulation of tau pathology in lower layers of both salience and executive-related regions of bvFTD patients.^21^ Other studies concentrated to cingulo-insular cortices associated with early salience-related symptoms in bvFTD show that select projection neurons comprising lower layers have similar vulnerabilities to both TDP-43 and tau pathology.^22–25^ Tau and TDP-43 pathology may target distinct populations of neurons in other regions, but patterns of neurodegeneration outside of salience-related regions in bvFTD remain understudied.

We address this gap in a large autopsy cohort of bvFTD by measuring immunoreactivity of neuronal markers in anterior cingulate, paracingulate, orbitofrontal, and middle frontal gyri due to their diverse functional, structural, and connectivity attributes. Functionally, these frontal regions serve key roles in one or more cognitive networks disrupted in bvFTD, including the anterior (para)cingulate in the salience network, the orbitofrontal cortex in the semantic-appraisal network, and the middle frontal cortex in the executive-control network.^2–7^ Structurally, these frontal regions have cytoarchitectonic subregions that are simultaneously distinct and spatially continuous due to their systematic transitions along the cortical mantle. This topologic arrangement of cytoarchitectonic subregions is known as a cortical gradient and in the frontal lobe includes phylogenetically older periallocortex and mesocortex in medial areas (e.g., cingulate gyri) that gradually transition into newer isocortex located in more lateral areas (e.g., orbitofrontal/middle frontal gyri).^26–28^ While antemortem neuroimaging lacks the resolution to discern gradual transitions between cytoarchitecture, histologic examinations can distinguish the spectrum of cytoarchitectonic types by the degree of laminar differentiation and predominance of pyramidal neurons in supragranular layers compared to infragranular layers.^26,27,29^ Thus, a postmortem parcellation of frontal regions presents a unique opportunity to investigate selective vulnerability and disease accumulation and spread between understudied cytoarchitectonic subregions. Indeed, not only do cytoarchitectonic types predict patterns of feedforward and feedback neuronal connectivity between layers of nearby and distant regions,^26,30–36^ changes in cortical layers may also influence connectivity in large-scale functional networks.^37–42^ Therefore, interconnected types of cytoarchitecture may provide a valuable anatomical model to conduct a novel comparative study of laminar neurodegeneration to identify the layer-specific microcircuits contributing to dysfunctional neurocognitive networks in bvFTD.

The aim of this study was to test the hypothesis that tauopathies, relative to TDP-43 proteinopathies, cause greater pyramidal neurodegeneration in clinically relevant frontal cytoarchitecture of bvFTD. We digitally quantified the immunoreactivity of total neurons and layer-specific pyramidal neurons in cytoarchitectonic subregions of the cortical gradient confirmed by our HC group and consistent with previous studies.^27,29,38,43^ Our main findings suggest that bvFTD-TDP and bvFTD-tau are clinically similar groups that show divergent patterns of vulnerability in cytoarchitecture along the cortical gradient, including pyramidal neurodegeneration that worsens along the mesocortical-to-isocortical gradient of cytoarchitecture in bvFTD-tau, but not bvFTD-TDP. Distinct cytoarchitectonic patterns of pyramidal neurodegeneration in the bvFTD spectrum may reflect cellular etiologies and patterns of spread that are proteinopathy-specific and informative to future biomarkers and therapeutics for clinical intervention.

## MATERIALS and METHODS

### Participants

Participants were clinically evaluated at the University of Pennsylvania (Penn) Frontotemporal Degeneration Center and clinical diagnoses were determined using modern consensus guidelines in weekly consensus meetings.^1,44–47^ Brain autopsies were performed at the Penn Center for Neurodegenerative Disease Research. As described previously,^48^ fresh tissue was sampled at autopsy and fixed overnight in 10% neutral buffered formalin or 70% ethanol with 150mmol NaCl. Neuropathologic diagnoses were made by expert neuropathologists using established criteria.^49–52^ All participants were prospectively genotyped for pathogenic mutations on FTD-associated genes based on a structured pedigree analysis described previously.^53^ All procedures in this study were performed in accordance with the standards of the Penn Institutional Review Board and the Declaration of Helsinki. Participant data were retrieved from the Penn Integrated Neurodegenerative Disease Database and retrospective chart review performed by experienced researchers (D.J.I., D.T.O.).^54^

We selected participants who met modern clinical criteria for bvFTD (n=77) and had a primary neuropathologic diagnosis of either FTLD-TDP (bvFTD-TDP) or FTLD-tau (bvFTD-tau). We excluded one bvFTD-TDP participant due to the presence of co-morbid cortical tauopathies (i.e., chronic traumatic encephalopathy, age-related tau astrogliopathy) that might obscure the interpretation of TDP43-mediated neurodegeneration. Additionally, one bvFTD-TDP and one bvFTD-tau participant were autopsied prior to modern clinical criteria and excluded due to limited chart data that precluded confirmation of bvFTD clinical diagnosis. We included a healthy control (HC) reference group comprised of those without a neurologic disorder/dementia and with low or no age-related pathology. After these exclusions, the three main groups of comparison included HC (n=32), bvFTD-TDP (n=47), and bvFTD-tau (n=27). Additional characteristics of participants meeting inclusion criteria are summarized in **Table 1**.

**Table 1.**
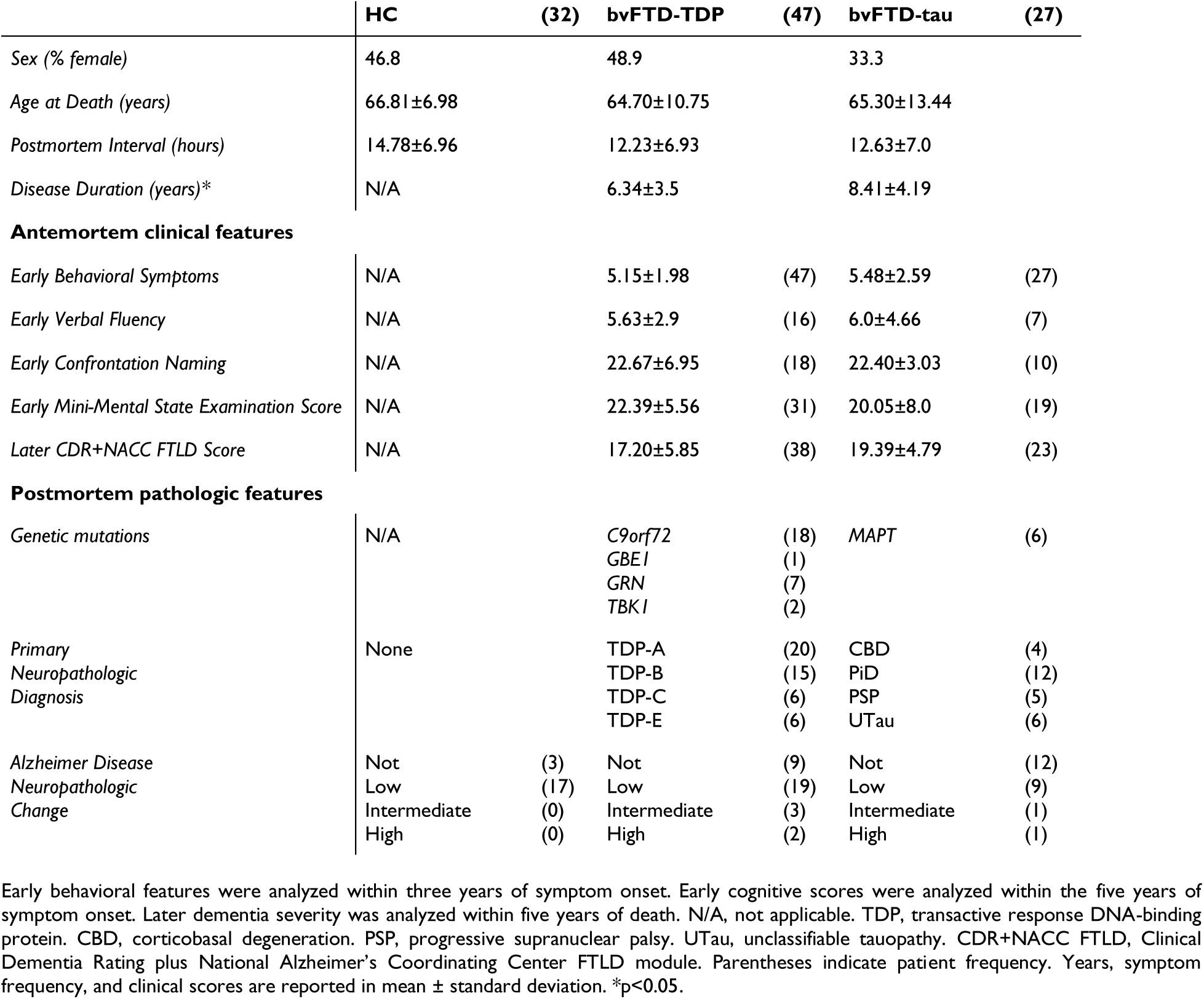
Patient demographics, clinical features, and pathologic features.

### Clinical data

Investigators D.J.I. and D.T.O. analyzed retrospective chart review data in all bvFTD participants to determine the frequency of early behavioral symptoms characteristic of bvFTD^18^ and performed retrospective Clinical Dementia Rating plus National Alzheimer’s Coordinating Center FTLD module (CDR+NACC FTLD)^55,56^ for each clinical visit in the record, as done previously.^57^ Presence and absence of behavioral symptoms within three years of symptom onset were evaluated by core diagnostic categories per clinical criteria.^1^ Behavioral symptoms were considered present if recorded at least once; symptoms were considered absent if not explicitly mentioned or could not be extrapolated from the records. All present symptoms were summed per participant, resulting in a maximum composite behavioral score ranging from 0–13. Within five years of symptom onset, we collected available neuropsychological data to evaluate cognitive impairment closest to clinical diagnosis, including letter fluency as a test of executive functioning,^58,59^ the Boston Naming Test for semantic retrieval,^60^ and Mini-Mental State Examination (MMSE) for global cognitive impairment. Lastly, we included CDR+NACC FTLD within five years of death to assess global dementia severity at later stages (**Table 1**).

### Immunohistochemistry and digital histopathology

We used a series of semi-adjacent paraffin-embedded 6µm-thick sections per main region (see Regions of interest below) to perform immunohistochemical procedures in the Penn Digital Neuropathology Lab as described previously.^21,61^ Immunohistochemistry for neuronal markers included antigen retrieval using citrate buffer and heat for neuronal nuclei protein (NeuN) enriched in most neurons (Anti-NeuN mouse monoclonal antibody, clone A60,1:1000, Millipore) and heavy non-phosphorylated neurofilament (NF-H) enriched in large pyramidal neurons (Anti-NF-H mouse monoclonal antibody, clone SMI32, 1:2000, Biolegend). All sections were counterstained with hematoxylin for further visualization of cell organization used in cytoarchitectonic type identification explained below and in **Supplementary Table 1**.

Immunostained sections were imaged on a digital slide scanner (Aperio AT2, Leica Biosystem, Wetzlar, Germany) at 20x magnification. Images were imported into QuPath software (version 0.2.0) to manually annotate cortical layers and subregions of interest described below. Within annotated areas we performed digital quantification similar to recent work,^19,21^ which included the use of an adaptive thresholding approach to digitally quantify the percent area occupied (%AO) by pixels with NeuN immunoreactivity (ir) and SMI32-ir (see details in **Supplementary Fig. 1**).

### Regions of interest

We measured NeuN-ir and SMI32-ir in three main regions of the frontal lobe from bilateral hemispheres where available, including anterior cingulate/paracingulate cortex (aCC), medial orbitofrontal cortex (mOFC), and middle frontal cortex (MFC) (coronal plane, **Fig. 1**). In the sagittal plane, MFC and mOFC regions were sampled approximately halfway along the anterior-posterior axis. The aCC, mOFC, and MFC collectively comprise up to six Brodmann areas (BA) and subregions reflecting at least five types of cytoarchitecture distinguishable by intrinsic cytoarchitectonic features (i.e., laminar organization, neuronal composition) and by their sequential positioning along gradients of laminar differentiation/expansion (i.e., “cortical gradients”).^27,29,62–66^ Cortical gradients reflect the topologic arrangement of cytoarchitectonic types as they gradually transition in an invariant allocortical-mesocortical-isocortical order, producing predictable topographic distributions.^26,28,29,43,67–69^ Moreover, cortical gradients are characterized by the progressive enlargement and density of pyramidal neurons in the external pyramidal layers II-III relative to the internal pyramidal layers V-VI, resulting in a range of “internopyramidal” and “externopyramidal” regions.^26,28,29,69^ In the current study, we sampled three types of mesocortex and two types of isocortex that belong to the parahippocampal and paraolfactory gradients found along medial-to-lateral axes of the dorsal and ventral frontal lobe.^26,28,29,68–70^ These five cytoarchitectonic types were numerated 1-5 in the current study due to their topologic order along the cortical gradient reflecting increasing externopyramidization. All topographic and cytoarchitectonic characteristics^62,71–73^ used for delineating cytoarchitectonic types are found in **Supplementary Table 1**.

**Figure 1.**
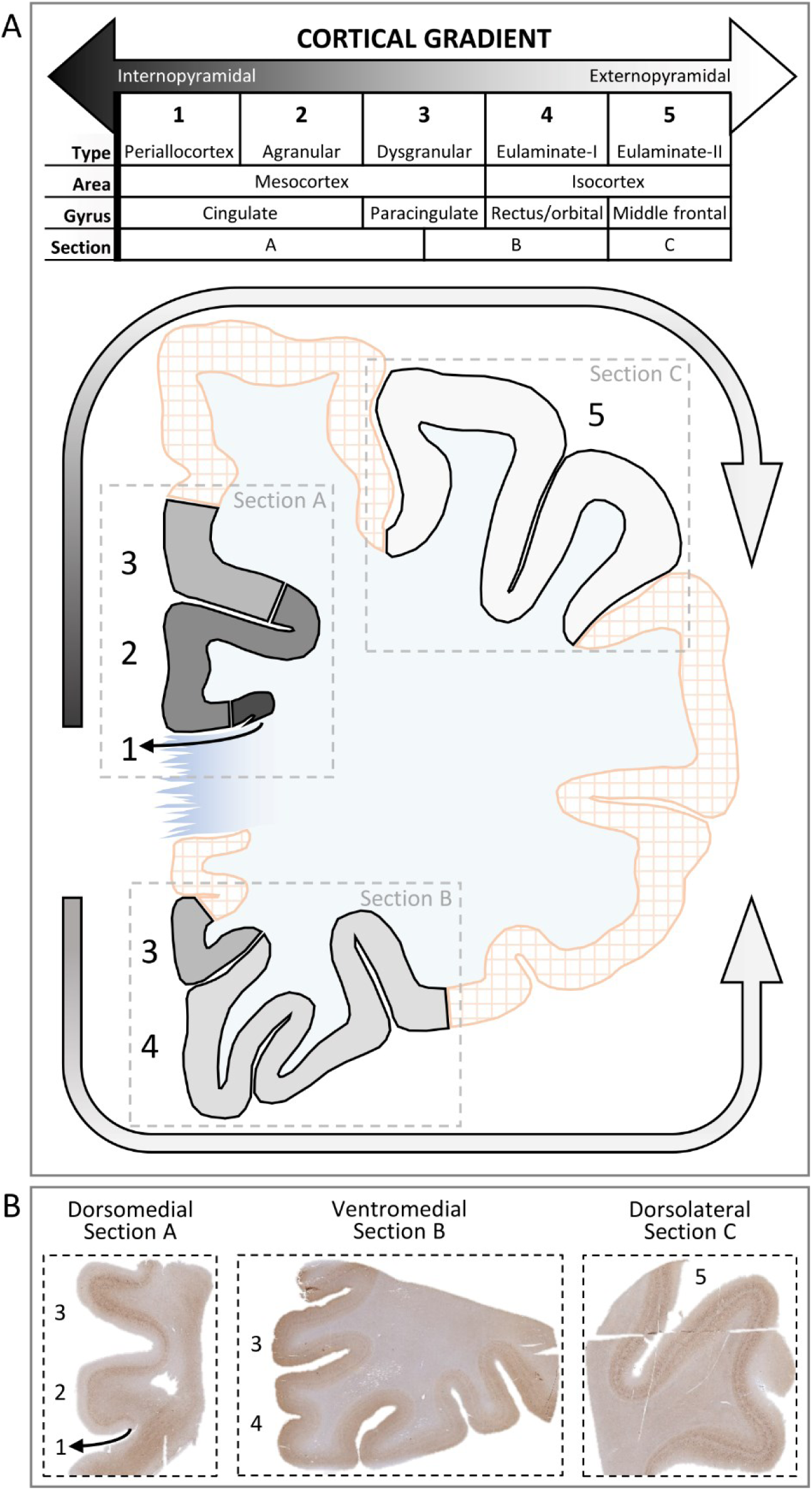
Regions and cytoarchitectonic subregions of interest along the cortical gradient of the frontal lobe. **A.** The frontal lobe cortical gradient is comprised of distinct subregions referred to as “cytoarchitectonic types” and numbered 1-5 due to their topological order. The predominance of supragranular pyramidal neurons progressively increases along the cortical gradient and defines internopyramidal from externopyramidal cytoarchitecture (depicted by gray scale arrows). Coronally, the cortical gradient transitions from internopyramidal cytoarchitecture in medial mesocortices (darker gray) to externopyramidal cytoarchitecture in more lateral isocortices (lighter gray). Cytoarchitectonic types have a predictable topography distributed across specific gyri found in postmortem tissue sections labeled A-C (delineated by dashed-line boxes). **B.** Representative HC tissue immunostained for SMI32 to demonstrate the three main tissue sections available per hemisphere (i.e., sections A-C) used to sample and quantify neurodegeneration in five types of cytoarchitecture #1-5.

In tissue immunostained stained for both NeuN and SMI32, a trained neuroanatomist (D.T.O.) first used topologic principles, topographic landmarks, and cytoarchitectonic features to reliably annotate subregions of each cytoarchitectonic type available per patient using a belt-transect method previously described.^21^ Next, all layers I-VI and pyramidal layers (i.e., supragranular II-III and infragranular V-VI) were separately annotated for digital measurements. All annotated subregions and layers closely corresponded to annotations made in semi-adjacent sections immunostained for NeuN and SMI32. Cytoarchitectonic types were not annotated due to tissue folds, tears, staining artifacts, or dissections excluded subregions of interest. A total of 1,024 distinct cytoarchitectonic types met inclusion criteria for laminar analyses of NeuN-ir and SMI32-ir in the current study (**Supplementary Table 2**).

### Staging levels of total neurodegeneration with NeuN-ir

To explore whether SMI32-ir loss is an earlier or later event in the neurodegenerative process, we assessed SMI32-ir in the context of total neurodegeneration measured by NeuN-ir. We used NeuN-ir to stage total neurodegeneration per cytoarchitectonic type because NeuN is a pan-neuronal marker sensitive to neurodegenerative processes^74^ including disease severity in the FTLD spectrum.^75,76^ Furthermore, we validated the digital quantitation of NeuN-ir by finding close correspondence between digital %AO by NeuN-ir and ordinal ratings of NeuN-ir completed in each layer from all available tissue in MFC (**Supplementary Fig. 2**). To stage total neurodegeneration in each cytoarchitectonic type, we included all available tissue from HC, bvFTD-TDP, and bvFTD-tau and calculated quartiles of NeuN-ir (i.e., sparse, low, intermediate, high). Thus, each NeuN-ir quartile reflects a level of total neurodegeneration (i.e., minimal, mild, moderate, or severe) that may indicate stages of early- to later-involved cortices, respectively (**Supplementary Table 3, Supplementary Fig. 3**). All cytoarchitectonic types with NeuN-ir were staged and used in analyses to determine if bvFTD groups showed different SMI32-ir between stages of total neurodegeneration.

### Statistical analyses

The %AO by SMI32-ir pyramidal neurons were natural log (ln) transformed to normalize data and then rescaled to 0-1 using min-max normalization for consistent interpretability in plots. To objectively determine the hierarchical position/rank of a cytoarchitectonic type along cortical gradients of “externopyramidization” in HC tissue, we measured the predominance of layer II-III pyramidal neurons (SMI32-ir^II-III^) using a calculation previously validated in healthy human and non-human cortical gradients^27,29,31,36,37,40,77,78^ as follows:

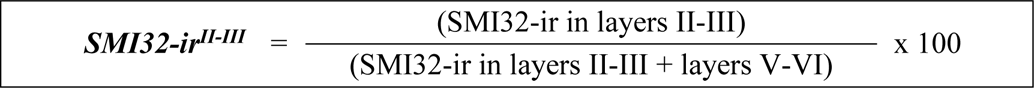

where smaller values represent more internopyramidal (mesocortical) areas of lower rank and larger values represent more externopyramidal (isocortical) areas of higher rank.

Group comparisons of demographics were analyzed using Chi-square analyses, *t*-test, or analysis of variance. We used linear regression adjusted for hemisphere, sex, ADNC, and age of death to determine the relationship between the ordinal variable cortical gradient (i.e., increasing laminar complexity of cytoarchitectonic types from 1-5) and the outcome measure SMI32-ir^II-III^ in the HC group. We analyzed neurodegeneration using linear mixed-effects (LME) models that included SMI32-ir (i.e., in all layers, supragranular layers, or infragranular layers) as the dependent variable and random intercepts for individual patients to account for correlations among repeated outcome measures from multiple regions. All LME models were adjusted for hemisphere, cytoarchitectonic type, sex, ADNC, and age at death. Disease duration was included as an additional covariate in models comparing SMI32-ir in each cytoarchitectonic type between bvFTD-TDP and bvFTD-tau. To determine the relationship between SMI32-ir and cognitive impairments, LME models included the covariates education and year interval between symptom onset and clinical evaluation. Finally, all LME models that analyzed group differences in SMI32-ir included NeuN-ir as a covariate to adjust for total neurodegeneration. Missing histologic data due to tissue unavailability or poor integrity were accounted for in all LME models and total group participants were reported in each result. Significance was set to *p*<0.05 and Bonferroni correction for multiple comparisons between groups was conducted. All analyses were performed using SPSS (version 29; Chicago, IL).

### Data availability

The datasets collected and/or analyzed during the current study are available from the corresponding author upon reasonable request.

## RESULTS

HC, bvFTD-TDP, and bvFTD-tau were similar in sex, age, and postmortem interval, but disease duration was longer in bvFTD-tau vs bvFTD-TDP (t[1,72]=2.273, p=0.026). Early behavioral severity, cognitive scores, and global dementia severity closer to death were all similar between bvFTD-TDP and bvFTD-tau (**Table 1, Supplementary Fig. 4**).

### Laminar distributions of SMI32-ir confirm the topologic position of distinct cytoarchitectonic types along the cortical gradient in HC

To determine that the five cytoarchitectonic types represent the continuous arrangement predicted by cortical gradients of the human brain, we tested whether the relative proportion of supragranular SMI332-ir neurons in HC tissue increased from internopyramidal mesocortex to more externopyramidal isocortex as has been shown previously (**Fig. 2A**).^27,29,38,43^ As expected in HC, we find that the cortical gradient was positively related to both the average SMI32-ir in all layers (β=0.013, SE=0.006, *p*=0.038; **Fig. 2B**) and %SMI32-ir^II-III^ (β=0.04, SE=0.006, *p*<0.001; **Fig. 2C**). Therefore, the cortical gradient of healthy cortex is characterized by an overall increase in SMI32-ir that transitions from infragranular-predominance (internopyramidal) to a more bilaminar/externopyramidal distribution (**Fig. 2-3**).

**Figure 2.**
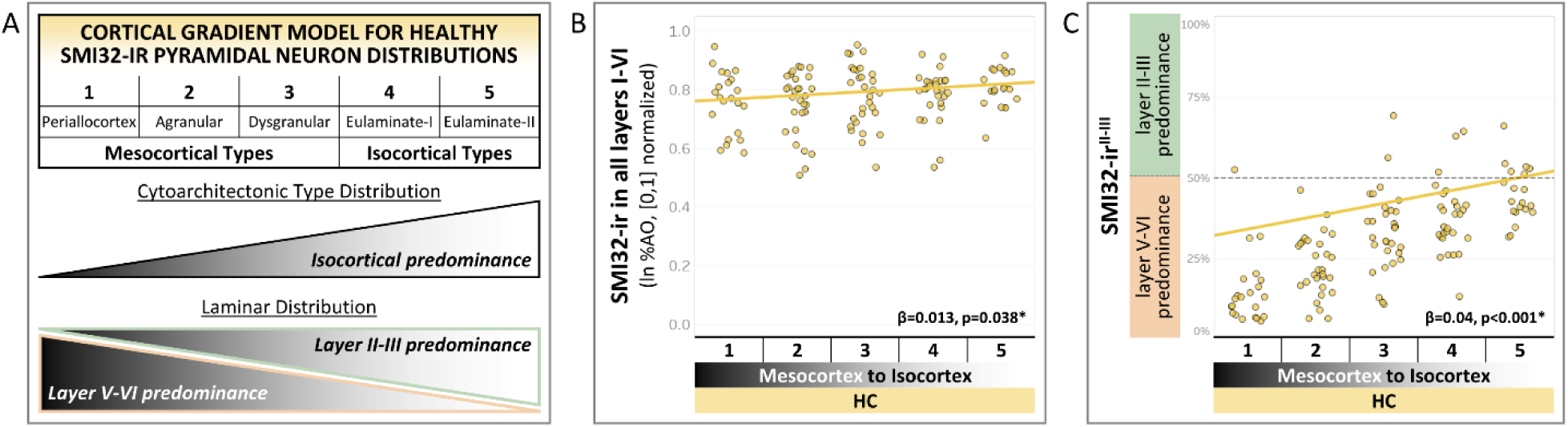
The cortical gradient predicts expected distributions of SMI32-ir in HC. **A.** Schematic diagram of expected distributions of SMI32-ir in HC tissue across the cortical gradient. **B.** SMI32-ir is positively related to the mesocortical-to-isocortical topologic arrangement of cytoarchitectonic types (β=0.013, SE=0.006, p=0.038), demonstrating that pyramidal neurons gradually become denser in isocortex relative to mesocortex. **C.** The SMI32-ir ratio of supragranular layers II-III to all pyramidal layers II,III,V,VI is related to the mesocortical-to-isocortical topologic arrangement of cytoarchitectonic types (β=0.04, SE=0.006, p<0.001), demonstrating our sampling approach includes relatively distinct cytoarchitectonic types found along the cortical gradient. Trend lines are derived from linear models adjusted for hemisphere, sex, ADNC stage, and age at death.

**Figure 3.**
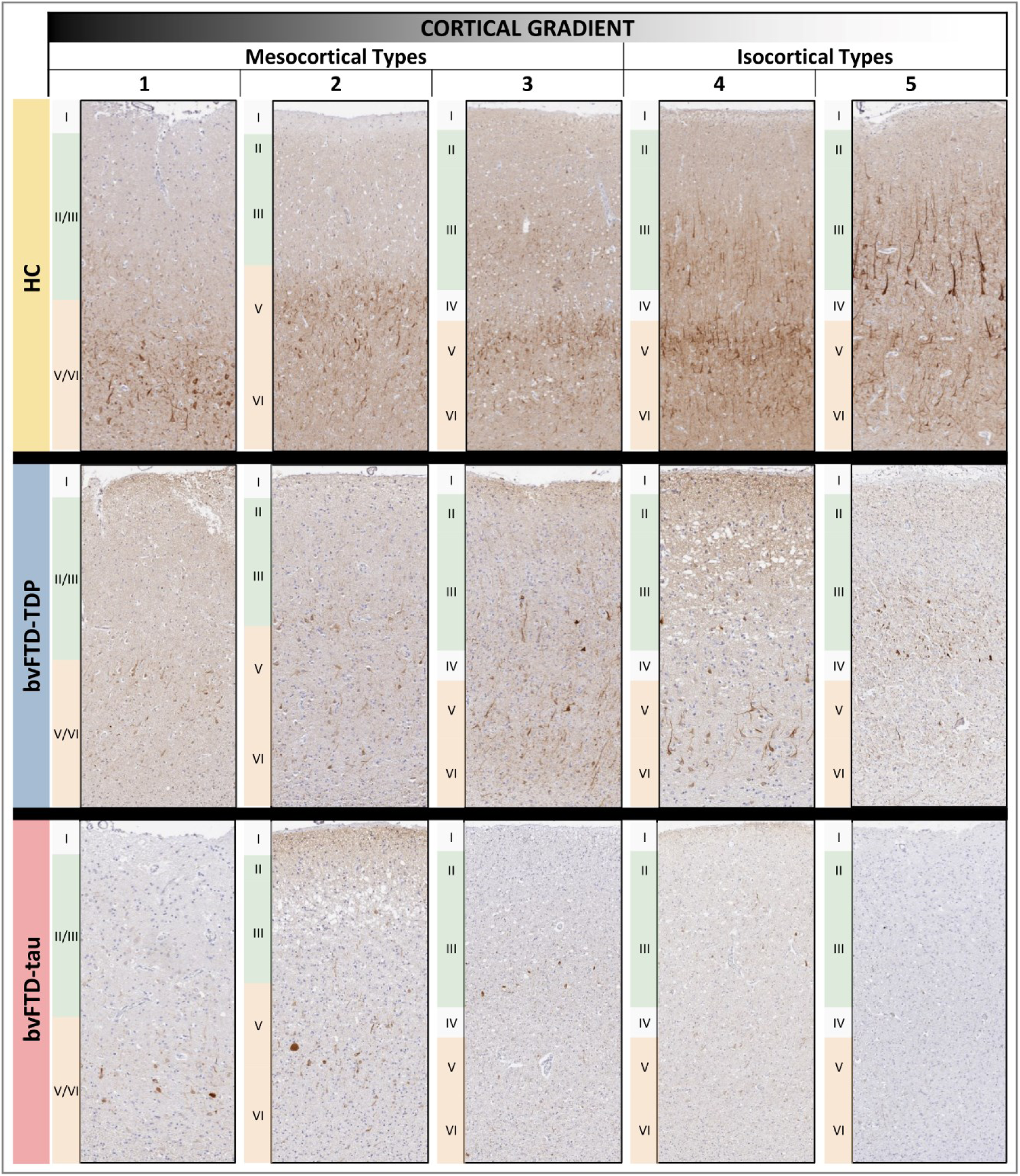
Representative photomicrographs depict distinct distributions of SMI32-ir pyramidal neurons along the cortical gradient in HC, bvFTD-TDP, and bvFTD-tau. As expected for HC tissue, SMI32-ir progressively increases from mesocortices to isocortices due to increasing size and density of predominantly layer III pyramidal neurons. In bvFTD-TDP, reduced levels of SMI32-ir appear largely similar between cytoarchitectonic types along the cortical gradient in bvFTD-TDP. In bvFTD-tau, there is a progressive reduction in SMI32-ir across the cortical gradient in bvFTD-tau that peaks in isocortex, especially type 5 (eulaminate-II) isocortex.

### Cortical gradient predicts divergent patterns of laminar pyramidal neurodegeneration between bvFTD-TDP and bvFTD-tau

We investigated whether the cortical gradient is related to distinct laminar distributions of SMI32-ir in each main group of HC, bvFTD-TDP, and bvFTD-tau (**Fig. 4A-B**). First, we found that SMI32-ir in supragranular layers II-III is positively related to the cortical gradient in HC (β=0.05, SE=0.006, *p*<0.001) and bvFTD-TDP (β=0.032, SE=0.005, *p*<0.001), but not related to the cortical gradient in bvFTD-tau (*p*=0.176), (**Fig. 4A**). Furthermore, a separate model testing the interaction term for group (i.e., bvFTD-TDP vs bvFTD-tau) and cortical gradient was significant (β=-0.024, SE=0.009, *p*=0.005), indicating that supragranular pyramidal neurodegeneration progressively worsens along the cortical gradient in bvFTD-tau relative to bvFTD-TDP.

**Figure 4.**
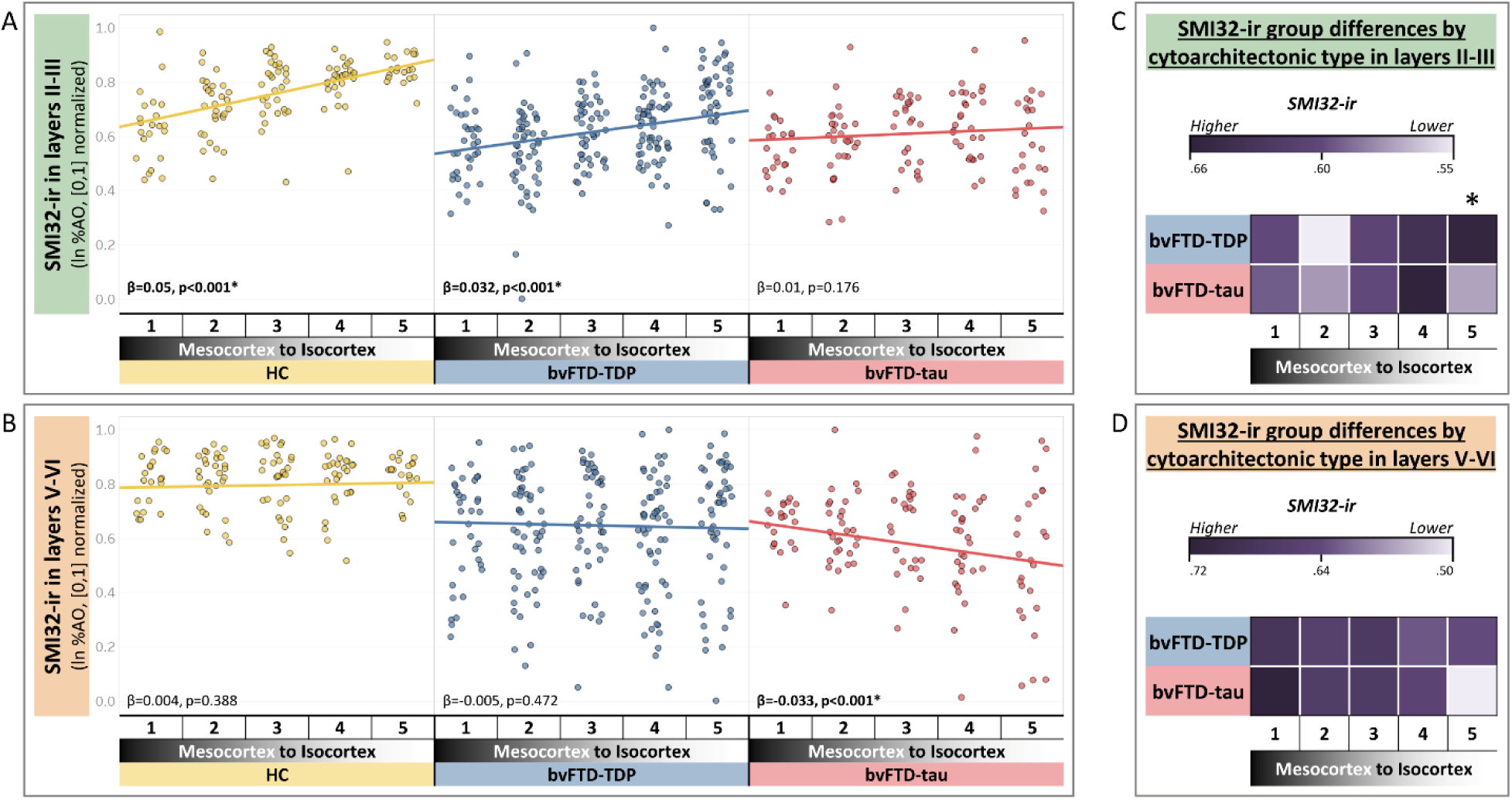
Cortical gradient predicts divergent patterns of laminar pyramidal neurodegeneration between bvFTD-TDP and bvFTD-tau. **A.** SMI32-ir in supragranular layers II-III is positively related to the cortical gradient in (β=0.05, SE=0.006, p<0.001) and bvFTD-TDP (β=0.032, SE=0.005, p<0.001), but not related to the cortical gradient in bvFTD-tau (p=0.176). Furthermore, a separate model testing the interaction term for group (i.e., bvFTD-TDP vs bvFTD-tau) and cortical gradient was significant (β=-0.024, SE=0.009, p=0.005), indicating that supragranular pyramidal neurodegeneration progressively worsens along the cortical gradient in bvFTD-tau relative to bvFTD-TDP. **B.** SMI32-ir in infragranular layers V-VI is not related to the cortical gradient in HC (p=0.388) or bvFTD-TDP (p=0.472), but negatively related to the cortical gradient in bvFTD-tau (β=-0.033, SE=0.009, p<0.001). Furthermore, a separate model testing the interaction term for group (i.e., bvFTD-TDP vs bvFTD-tau) and cortical gradient was significant (β=-0.03, SE=0.011, p=0.004), indicating that infragranular pyramidal neurodegeneration progressively worsens along the cortical gradient in bvFTD-tau relative to bvFTD-TDP. Trend lines are derived from LME models adjusted for hemisphere, sex, ADNC stage, age at death, and NeuN-ir in all layers. Heat maps show the quantitative distribution of SMI32-ir in layers II-III **(C)**, layers V-VI **(D)** of each cytoarchitectonic type in bvFTD-TDP and bvFTD-tau. Shades of purple represent estimated means of normalized ln SMI-32-ir in each region derived from linear mixed-effects models that compared groups and adjusted for hemisphere, sex, ADNC stage, age at death, disease duration, and NeuN-ir in combined layers. We find that both supragranular and infragranular SMI32-ir is similar between bvFTD-TDP and bvFTD-tau in all cytoarchitectonic types except in type 5 where we find supragranular SMI32-ir is lower in bvFTD-tau vs bvFTD-TDP (β=-0.086, SE=0.041, p=0.039*).

Next, we found that SMI32-ir in infragranular layers V-VI is not related to the cortical gradient in HC (*p*=0.388) or bvFTD-TDP (*p*=0.472), but negatively related to the cortical gradient in bvFTD-tau (β=-0.033, SE=0.009, *p*<0.001), (**Fig. 4B**). Furthermore, a separate model testing the interaction term for group (i.e., bvFTD-TDP vs bvFTD-tau) and cortical gradient was significant (β=-0.03, SE=0.011, *p*=0.004), indicating that infragranular pyramidal neurodegeneration progressively worsens along the cortical gradient in bvFTD-tau relative to bvFTD-TDP. We found similar relationships between the cortical gradient and laminar SMI32-ir for pathologic subgroups of bvFTD-TDP and bvFTD-tau (**Supplementary Fig. 5**). Moreover, similar results were found in models that excluded the covariate NeuN-ir (data not shown), suggesting that relationships between the cortical gradient and SMI32-ir are not significantly influenced by the variance in total neurodegeneration by cytoarchitectonic type.

Lastly, we compared laminar SMI32-ir between bvFTD-TDP and bvFTD-tau by cytoarchitectonic type (**Fig. 4C-D**). We find that both supragranular and infragranular SMI32-ir is similar between bvFTD-TDP and bvFTD-tau in all cytoarchitectonic types except in type 5 where we find supragranular SMI32-ir is lower in bvFTD-tau vs bvFTD-TDP (β=-0.086, SE=0.041, *p*=0.039), (**Fig. 4C**). Therefore, maximal pyramidal neurodegeneration varies along the cortical gradient between bvFTD-TDP and bvFTD-tau, with peak loss in supragranular layers of eulaminate-II isocortex of bvFTD-tau.

### Differential involvement of layers and cytoarchitecture connected by long-range circuits in bvFTD-tau vs bvFTD-TDP

According to the structural model of laminar connectivity,^26,30,31,35^ different types of cytoarchitecture are connected by predictable sets of layers along cortical gradients. Of the many pathways that interconnect the cortical gradient examined in our study, the longest pathways may connect infragranular layers of agranular mesocortex (cytoarchitectonic type 2) with supragranular layers of eulaminate-II isocortex (cytoarchitectonic type 5), (**Fig. 5A**).^79^ Thus, we used this laminar connectivity model to test the hypothesis that laminar SMI32-ir in these select areas would show differential involvement between bvFTD-tau and bvFTD-TDP.

**Figure 5.**
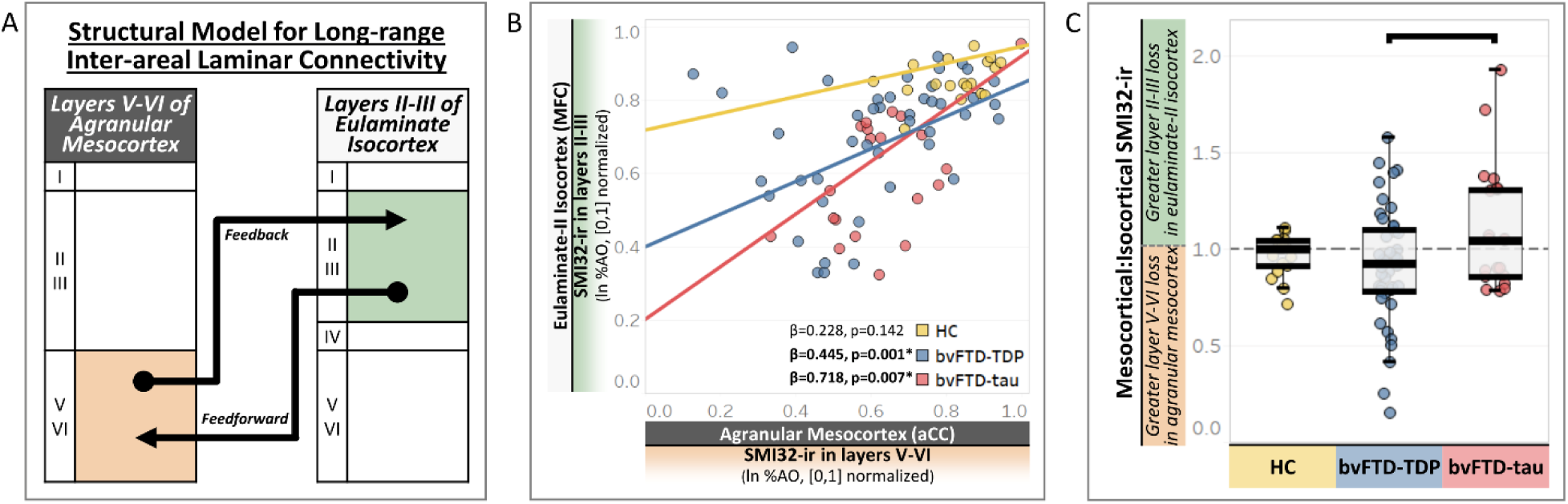
Layers and cortices connected by longer-range circuits show greater pyramidal neurodegeneration in bvFTD-tau vs bvFTD-TDP. **A.** A schematic depicting the ‘structural model’ of neuronal connectivity in the cortical gradient whereby a subset of long-range pathways comprising feedback and feedforward circuits connect dissimilar cytoarchitecture types (e.g., agranular mesocortex and eulaminate isocortex) via supragranular layers V-VI in mesocortex and infragranular layers II-III in isocortex. **B.** We find that lower SMI32-ir in infragranular mesocortex predicted lower SMI32-ir in supragranular isocortex in both bvFTD-TDP (n=45 patients; β=0.445, SE=0.098, p=0.001) and bvFTD-tau (n=25 patients; β=0.718, SE=0.227, p=0.007). In contrast, we find no significant association between SMI32-ir in infragranular mesocortex and supragranular eulaminate-II isocortex in HC (n=18 patients; p=0.142), reflecting the consistently high SMI32-ir between interconnected layers. **C.** In an exploratory analysis of infragranular mesocortex-to-supragranular isocortex ratios of SMI32-ir, we find significantly larger ratios in bvFTD-tau compared to bvFTD-TDP (β=0.188, SE=0.078, p=0.019), suggesting preferential supragranular isocortical neurodegeneration in bvFTD-tau compared to preferential infragranular mesocortical neurodegeneration in bvFTD-TDP.

In support of the structural model and the influence laminar connectivity may have on neurodegeneration in bvFTD, we stratified analyses by group to find that lower SMI32-ir in infragranular mesocortex predicted lower SMI32-ir in supragranular isocortex in both bvFTD-TDP (n=45 patients; β=0.445, SE=0.098, *p*=0.001) and bvFTD-tau (n=25 patients; β=0.718, SE=0.227, *p*=0.007) which graphically appeared to have a stronger correlation in bvFTD-tau, but not in HC (n=25 patients; *p*=0.142), (**Fig. 5B**). Next, we performed a planned comparison between bvFTD-TDP and bvFTD-tau in an exploratory LME model using SMI32-ir ratios of infragranular mesocortex-to-supragranular isocortex to test for differential involvement between the distant interconnected layers. We find significantly larger ratios in bvFTD-tau compared to bvFTD-TDP (β=0.188, SE=0.078, *p*=0.019), (**Fig. 5C**), suggesting preferential supragranular isocortical neurodegeneration in bvFTD-tau compared to preferential infragranular mesocortical neurodegeneration in bvFTD-TDP. Moreover, these data provide converging evidence that select long-range laminar microcircuits known to link distant and distinct cortical types along the cortical gradient may influence the severity of neurodegeneration between subregions, including greater isocortical pyramidal neurodegeneration in bvFTD-tau compared to bvFTD-TDP.

### Evidence of earlier pyramidal neurodegeneration in bvFTD-tau vs bvFTD-TDP

To test the hypothesis that distinct proteinopathies may cause pyramidal neurons to degenerate disproportionately early where overall neurodegeneration is minimal, we performed exploratory analyses of SMI32-ir in subsets of tissue with sparse/low NeuN-ir, intermediate NeuN-ir, or high NeuN-ir (**Fig. 6A-B**).

**Figure 6.**
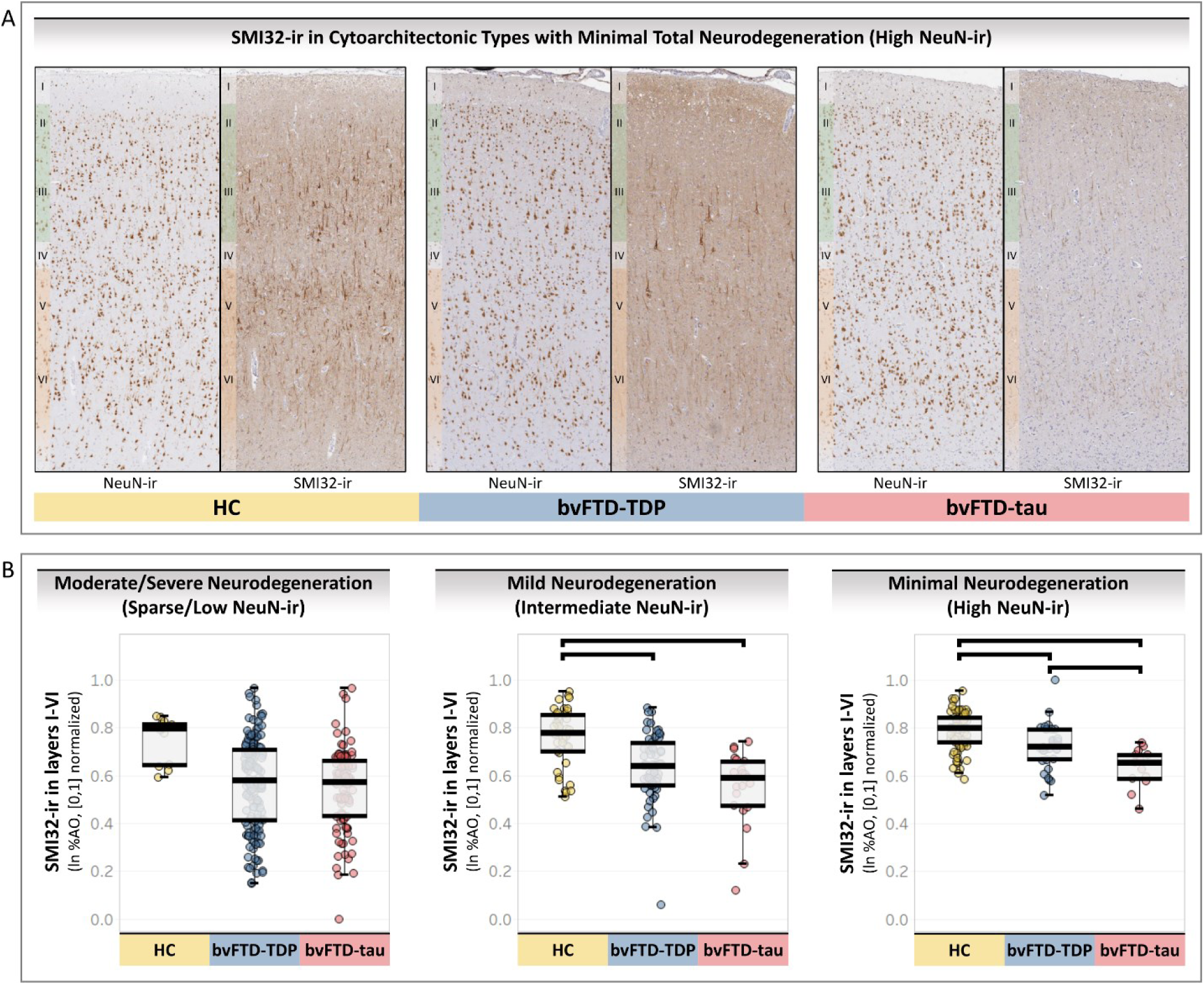
Evidence of earlier pyramidal neurodegeneration in bvFTD-tau vs bvFTD-TDP. **A.** Representative microphotographs of cytoarchitecture with high NeuN-ir in comparison to varied levels of SMI32-ir in semi-adjacent tissue by group. Despite high preservation of NeuN-ir neurons, note the prominent reduction in SMI32-ir pyramidal neurons in bvFTD-tau relative to bvFTD-TDP and HC tissue. **B.** In cytoarchitectonic types with sparse-to-low NeuN-ir, we find no significant differences in SMI32-ir between HC, bvFTD-TDP, and bvFTD-tau (F[2,72.3]=1.316, p=0.275). In cytoarchitectonic types with intermediate NeuN-ir, we find lower SMI32-ir in bvFTD-tau (β=-0.199, SE=0.048, p<0.001) and bvFTD-TDP (β=-0.143, SE=0.039, p=0.002) compared to HC, while SMI32-ir is similar between bvFTD-TDP and bvFTD-tau (p=0.713). In cytoarchitectonic types with the highest NeuN-ir, we find lower SMI32-ir in bvFTD-tau (β=-0.187, SE=0.036, p<0.001) and bvFTD-TDP (β=-0.076, SE=0.03, p=0.046) compared to HC. We additionally find lower SMI32-ir in bvFTD-tau compared to bvFTD-TDP (β=-0.11, SE=0.044, p=0.047).

In an LME model that only included cytoarchitectonic types with sparse-to-low NeuN-ir, we find no significant differences in SMI32-ir between HC (n=7 patients), bvFTD-TDP (n=33 patients), and bvFTD-tau (n=23 patients) (F[2,72.3]=1.316, p=0.275).

In an LME model that only included cytoarchitectonic types with intermediate NeuN-ir, we find lower SMI32-ir in bvFTD-tau (n=10 patients; β=-0.199, SE=0.048, *p*<0.001) and bvFTD-TDP (n=24 patients; β=-0.143, SE=0.039, *p*=0.002) compared to HC (n=18 patients), while SMI32-ir is similar between bvFTD-TDP and bvFTD-tau (p=0.713).

In an LME model that only included cytoarchitectonic types with the highest NeuN-ir, we find lower SMI32-ir in bvFTD-tau (n=6 patients; β=-0.187, SE=0.036, *p*<0.001) and bvFTD-TDP (n=11 patients; β=-0.076, SE=0.03, *p*=0.046) compared to HC (n=26 patients). Interestingly, we additionally find lower SMI32-ir in bvFTD-tau compared to bvFTD-TDP (β=-0.11, SE=0.044, *p*=0.047). Therefore, significantly lower SMI32-ir in cytoarchitectonic types with high NeuN-ir/minimal overall neurodegeneration in bvFTD-tau suggests that pyramidal neurodegeneration may occur earlier or more severely in select regions and cases of bvFTD-tau compared to bvFTD-TDP and HC.

### Pyramidal neurodegeneration is related to the severity of early behavioral and cognitive impairment in bvFTD

Because mesocortex is more directly involved with behavioral changes in bvFTD,^80,81^ we stratified our analyses for mesocortical and isocortical types to determine if changes in behavior is associated with SMI32-ir in select areas. As expected for behavior-related mesocortex, we find that lower SMI32-ir in all layers I-VI is related to more behavioral symptoms (β=-0.018, SE=0.009, *p*=0.044), but no relationship is found in isocortex (p=0.458), (**Fig. 7A**).

**Figure 7.**
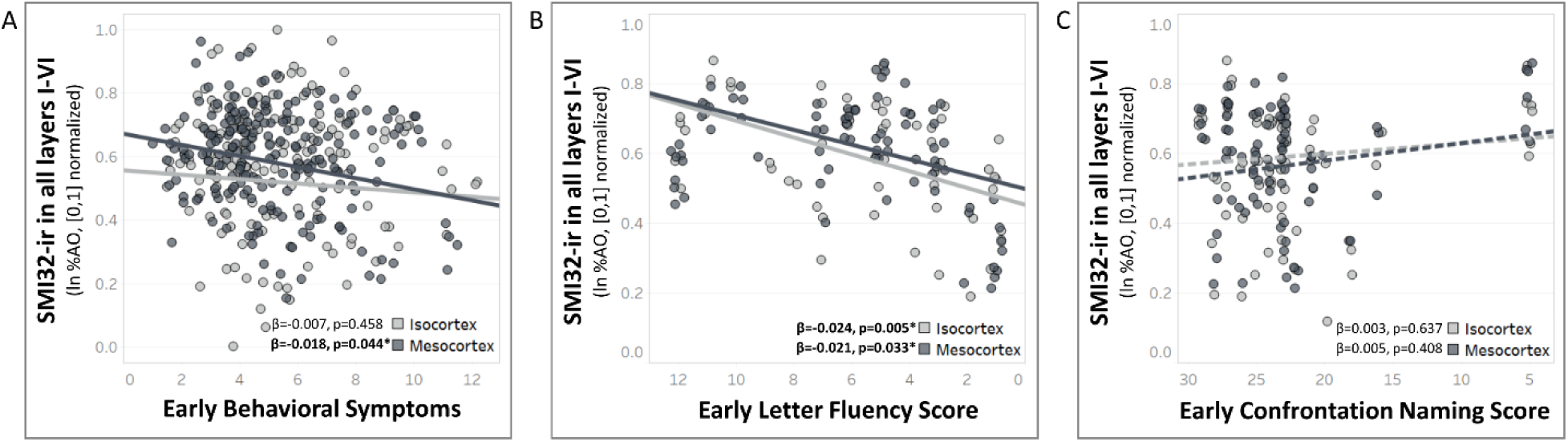
Relationships between pyramidal neurodegeneration and clinical features of bvFTD. **A.** As expected for paralimbic structures, more behavioral symptoms were associated with lower SMI32-ir in mesocortical types (i.e., periallocortex, agranular mesocortex, dysgranular mesocortex) (β=-0.018, SE=0.009, p=0.044), not isocortical types (i.e., eulaminate I-II). **B.** Worse performance on letter fluency was related to lower SMI32-ir in both mesocortex (β=0.021, SE=0.009, p=0.033) and isocortex (β=0.024, SE=0.007, p=0.005). **C.** Performance on confrontation naming was not related to SMI32-ir in either mesocortex or isocortex. Trend lines are derived from LME models adjusted for hemisphere, cytoarchitectonic type, sex, ADNC stage, age at death, bvFTD group, education, and interval between test date and death.

In a subset of patients with cognitive impairments evaluated close to symptom onset, we tested the hypothesis that pyramidal neurodegeneration is related to letter fluency due to its greater association with frontal lobe integrity^58^ than confrontation naming, a test that relies more so on temporal lobe integrity.^82^ We find that worse letter fluency performance is related to lower SMI32-ir in both mesocortex (n=23 patients; β=0.021, SE=0.009, *p*=0.033) and isocortex (n=23 patients; β=0.024, SE=0.007, *p*=0.005), (**Fig. 7B**). In contrast, we find that confrontation naming performance is not related to SMI32-ir in either mesocortex or isocortex (n=28 patients; p<0.05), (**Fig. 7C**), affirming the clinical relevance of SMI32-ir loss in frontal cytoarchitecture of bvFTD.

## DISCUSSION

Our comparative histopathologic study investigated proteinopathy-specific patterns of neurodegeneration in a pathologically heterogenous cohort of bvFTD patients. We hypothesized that bvFTD-tau shows greater pyramidal neurodegeneration than bvFTD-TDP within a cortical gradient framework. Because pyramidal neuron density intrinsically varies by layers and regions in healthy brains, we digitally quantified the area occupied by SMI32-ir pyramidal neurons across a mesocortex-to-isocortex gradient of cytoarchitectonic subregions in the frontal lobe of HC, bvFTD-TDP, and bvFTD-tau participants. First, we used digital measures of laminar SMI32-ir to recapitulate the cortical gradient in HC tissue. Next, we found that the cortical gradient predicts divergent relationships with pyramidal neurodegeneration in bvFTD-TDP versus bvFTD-tau. Using a model for laminar connectivity between relatively distant cytoarchitectonic types along the cortical gradient, we found that infragranular mesocortex predicted greater pyramidal neurodegeneration in supragranular eulaminate-II isocortex of bvFTD-tau compared to bvFTD-TDP. We show pyramidal neurodegeneration may be an earlier event in bvFTD-tau given that tissue with high levels of NeuN-ir neurons showed greater pyramidal neurodegeneration in bvFTD-tau compared to bvFTD-TDP. Finally, we find that early behavioral severity is related to greater pyramidal neurodegeneration in paralimbic mesocortices, not heteromodal isocortices.

In our retrospective clinical review, we found a similar prevalence of core diagnostic symptoms during the first three years of clinical progression in bvFTD-tau vs bvFTD-TDP (**Table 1**). BvFTD is a clinically heterogenous syndrome and others have found subtle differences in clinical features between molecular subtypes of FTLD.^18,83,84^ Despite the size of our rare autopsy cohort, we are limited in the ability to compare clinical data amongst pathologic subtypes of FTLD-tau and FTLD-TDP. Therefore, we cannot exclude that in larger numbers of patients, subtypes may have neuroanatomic differences in neurodegeneration that contribute to clinical heterogeneity in bvFTD. Importantly, we find our groups shared a similar severity of global (i.e., MMSE and CDR+NACC FTLD) and specific cognitive impairment, including letter fluency and confrontation naming (**Table 1**). Thus, our analyses highlight partially non-overlapping patterns of neurodegeneration in bvFTD-TDP vs bvFTD-tau despite similar clinical presentation and overall disease severity near autopsy.

### PYRAMIDAL NEURODEGENERATION INCREASES ALONG THE CORTICAL GRADIENT OF BVFTD-TAU, NOT BVFTD-TDP

To test our main hypothesis that bvFTD-tau was associated with greater pyramidal neurodegeneration, we examined patterns of neuron loss in the context of a well-known cortical gradient that we confirmed in our HC group. The frontal lobe cortical gradient is a neuroanatomical framework to reliably examine heterogeneous cytoarchitecture that includes allocortical, mesocortical, and isocortical types. Importantly, the allocortical-to-isocortical gradient is topologically arranged and has increasing densities of neurons^43,85^ with distinct laminar distributions of pyramidal neurons.^27,29,31,37,77,78^ In addition to the structural organization of cortical gradients, a growing literature suggests that they are evolutionary conserved across mammals^27,28,43,68,69,86^ and correspond to gradients of gene expression,^38,87,88^ distributions of neurotransmitters,^89^ patterns of circuit connectivity,^42,90,91^ and hierarchies of functional and cognitive networks.^92–97^ Therefore, measuring deviations to an otherwise healthy cortical gradient in neurodegenerative disorders can potentially inform why, and to what degree, pathologic changes have occurred where they were measured. Consistent with previous studies of healthy human cortex,^29,43^ we confirmed the reliable sampling of cytoarchitectonic types along the cortical gradient by demonstrating that the predominance of supragranular layer SMI32-ir pyramidal neurons increases along the cortical gradient of our healthy control cohort (**Fig. 2-3**). Within this anatomical framework, we found that pyramidal neurodegeneration progressively worsened along the cortical gradient in bvFTD-tau, not bvFTD-TDP (**Fig. 3-4**). Moreover, bvFTD-tau showed greater pyramidal loss in the supragranular layers of eulaminate-II isocortex relative to bvFTD-TDP that was not mediated by disease duration controlled for in our models. Therefore, tau-mediated degeneration in bvFTD may have a predilection for pyramidal neurons that steadily transition to supragranular-predominance in more isocortical areas of the cortical gradient as we previously observed.^19^ It is interesting to note that neurofibrillary tau pathology in AD has a similar selectivity for SMI32-ir neurons concentrated to layers III and V.^98–102^ Furthermore, single nucleus RNA-seq data and genome-wide association studies have identified genes that regulate tau homeostasis and autophagy are more frequent in the most vulnerable excitatory neurons in AD human brains and animal models.^103,104^ SMI32-ir demonstrates that pyramidal neurons are enriched for non-phosphorylated neurofilament, a cytoskeletal protein that can undergo abnormal hyperphosphorylation that leads to tau pathology. Therefore, one potential mechanism that links tauopathies and selective pyramidal neurodegeneration is that pyramidal neurons may represent a subpopulation more dependent on biochemical regulation of neurofilament and thus more vulnerable to dysregulation and degeneration by progressive tau aggregation. Importantly, while bvFTD-TDP showed SMI32-ir loss, it was less severe than SMI32-loss in bvFTD-tau and did not follow the cortical gradient. Future work will examine the anatomic principles that guide selective bvFTD-TDP neurodegeneration.

### LAYERS CONNECTING LONG-RANGE PATHS REVEAL DIFFERENTIAL VULNERABILITIES BETWEEN BVFTD-TDP AND BVFTD-TAU

Recent neuroimaging studies demonstrate that many lateralized isocortices are atrophied early and severely in bvFTD, suggesting that bvFTD has more of a multi-network profile of degeneration than a singular salience network-centric profile.^7^ Postmortem patterns of neurodegeneration outside salience-related mesocortices are understudied in bvFTD, but our investigation provides new insight into the neuronal vulnerability of eulaminate I & II isocortex found in orbitofrontal and middle frontal gyri. While bvFTD-TDP and bvFTD-tau showed significant pyramidal neurodegeneration relative to HC, our data suggests that bvFTD-tau has more severe neurodegeneration toward the isocortical end of the cortical gradient that peaks in the dorsolateral eulaminate-II isocortex (**Fig. 5**). As hypothesized earlier, one explanation for this pattern could be that tau-mediated degeneration has a greater propensity for neurofilament-rich projection neurons than TDP43-mediated degeneration. Moreover, pyramidal neurons are the origin of long-range corticocortical circuits that interconnect the cortical gradient, representing a structural determinant of where and how extensive neurodegeneration is in bvFTD-tau vs bvFTD-TDP. As opposed to the ubiquitous short-range pathways within and between layers of the same or similar cytoarchitectonic types, long-range pathways between distant cytoarchitectonic types form a signature laminar profile of connectivity whereby select sets of layers are more interconnected than others. Specifically, tract tracing studies and network models of neural connectivity suggest that feedback/feedforward circuits that connect the farther ends of cortical gradients are predominantly mediated by infragranular layers V-VI of mesocortex and supragranular layers II-III layers of isocortex (**Fig. 5A**).^26,30,32,35,36,79,105^ If neurodegeneration in these sets of layers is more pronounced in one group of bvFTD patients, it may reflect that this long-range pathway is selectively vulnerable to a distinct proteinopathy. Our current study modeled these long-range pathways via the laminar subregions of the cortical gradient and found that infragranular SMI32-ir in agranular mesocortex predicts supragranular SMI32-ir in eulaminate-II isocortex, which is significantly lower in bvFTD-tau compared to bvFTD-TDP (**Fig. 4-5**). Thus, longer-range circuits that connect mesocortex to isocortex may directly contribute to the elevated isocortical pyramidal neurodegeneration we observed in bvFTD-tau. These circuits may have particular clinical relevance given that a recent neuroimaging study in FTD found a closer link between long-range white matter tracts and executive dysfunction than shorter-range tracts.^106^ Neuroimaging studies and network modeling have implicated long-range neuronal pathways in FTLD-tau previously,^107,108^ in addition to histologic studies examining large-scale networks that connect cortex to brainstem and spinal structures. For example, we found a predilection for tau-mediated degeneration of noradrenergic neurons in the locus coeruleus compared to TDP-43 pathology in FTLD,^109^ and tau inclusions have been shown to preferentially spread along corticospinal and corticopontine fibers in FTLD.^110^ Therefore, bvFTD-tau may preferentially target long-range circuits and large networks that interconnect corticocortical and cortico-subcortical systems than bvFTD-TDP.

### EVIDENCE FOR EARLIER PYRAMIDAL NEURODEGENERATION IN BVFTD-TAU

Previous studies in relatively small cohorts of either FTLD-TDP or FTLD-tau have found pronounced degeneration of spindle (i.e. von Economo) neurons and fork neurons in anterior cingulate and insula mesocortices.^23,24,111,112^ Because these mesocortices are part of the salience network that might be the earliest involved in bvFTD,^2,3,6^ these subpopulations of infragranular projection neurons have been speculated to be the site of genesis and propagation of proteinopathies in bvFTD. A recent comparative study has taken this further by showing greater loss of infragranular projection neurons in the anterior cingulate of bvFTD patients with TDP-43 or FUS pathology compared to bvFTD-tau patients.^113^ However, our analysis of the same spindle neuron-rich agranular mesocortex found comparable levels of pyramidal neurodegeneration between bvFTD-TDP and bvFTD-tau, regardless of layer examined (**Fig. 3-5**). Furthermore, bvFTD-TDP and bvFTD-tau showed similar levels of SMI32-ir neurons in neighboring cingulate cytoarchitecture (i.e., periallocortex and dysgranular mesocortex), suggesting that frontal mesocortices are broadly vulnerable to pyramidal degeneration in the bvFTD spectrum. Our unique finding may be partly explained by our larger bvFTD cohort and more exhaustive sampling of the full cytoarchitectonic range of cingulate mesocortex, resulting in a more representative assessment of projection neuron loss compared to previous experimental designs. Another difference is our digital measure and SMI32 marker for long-projecting pyramidal neurons compared to the densities and markers for projection neurons used previously (e.g., GABRQ or morphologic classification via Nissl cresyl violet).^23,24,111–113^

Cingulate mesocortex and the salience network may not always be the earliest regions involved in bvFTD,^7^ but it is unknown if distinct proteinopathies target earlier involved regions differently. Despite the cross-sectional postmortem nature of histologic studies, cytoarchitectonic types with minimal overall neurodegeneration may be considered relatively earlier involved in the degeneration process. We used NeuN-ir as a proxy for overall neurodegeneration to test the hypothesis that bvFTD-tau would have greater pyramidal neurodegeneration in cytoarchitectonic types with high NeuN-ir compared to cytoarchitectonic types with lower levels of NeuN-ir (**Fig. 6**). In support of our hypothesis, only the cytoarchitectonic types with the highest NeuN-ir showed greater pyramidal neurodegeneration in bvFTD-tau than bvFTD-TDP. The cytoarchitectonic types with the highest NeuN-ir spanned all cytoarchitectonic types and pathologic subgroups, suggesting that pyramidal neurodegeneration may be potentially earlier in bvFTD-tau and involve a network of likely interconnected cortices of multiple structural types and function.

### PYRAMIDAL NEURODEGNERATION IS RELATED TO CLINICAL SEVERITY

We performed a series of clinicoanatomical analyses to determine the clinical relevance of SMI32-ir loss in distinct cytoarchitecture. Specifically, we determined that more frequent behavioral symptoms in our clinically similar cohort of bvFTD predicted lower SMI32-ir in mesocortices, not isocortices (**Fig. 7A**). While these findings are consistent with the clinical correlates of agranular cingulate mesocortex measured in previous postmortem studies of bvFTD,^22,24,112,114^ our inclusion of periallocortex and dysgranular mesocortex expands our understanding of the cingulate mesocortical structures that likely drive complex behavioral changes in bvFTD. For instance, dysgranular mesocortex of paracingulate gyri anatomically corresponds to nodes of salience and default mode networks, suggesting a role in core bvFTD-related symptoms such as apathy, empathy loss, and changes in hedonic values.^7,115,116^ The functions of cingulate periallocortex are poorly understood, but its phylogenetically old structure may subserve limbic processing of pain or emotion similar to neighboring cytoarchitecture.^73,117,118^ In contrast, phylogenetically newer structures in our investigation (i.e., eulaminate-I-II isocortices) are heteromodal association cortices more often associated with semantic appraisal or executive dysfunction than behavioral symptoms.^1,58,119,120^ In fact, we found that greater pyramidal neurodegeneration predicted worse performance on the letter fluency test for executive functioning, not confrontation naming (**Fig. 7B-C**). This is in line with previous antemortem neuroimaging studies that showed selective atrophy of anterior cingulate, orbitofrontal, and dorsolateral prefrontal cortices was associated with letter fluency in bvFTD.^58^ Moreover, our data support the conclusion that pyramidal neurons and their progressive loss in meso- and isocortical areas likely contribute to a combination of socioemotional deficits and impaired mental search strategies characteristic of bvFTD.

### PYRAMIDAL NEURODEGENERATION BETWEEN BVFTD SUBGROUPS

Our bvFTD cohort included a wide range of pathologic and mutation carrier subgroups to determine the consistency of pyramidal neurodegeneration across the bvFTD spectrum. In supplementary analyses (**Supplementary Fig. 5-6**) that compared all pathologic subgroups, we found more pronounced pyramidal neurodegeneration in PiD and PSP. In a separate analysis of mutation carriers and sporadic subgroups vs HC, we found that *C9orf72*, *GRN*, and sporadic bvFTD-tau subgroups showed more significant pyramidal neurodegeneration. These findings suggest that select subgroups show subtly more severe profiles of neuron loss consistent with previous histologic studies of PiD, *C9orf72*, and *GRN* in bvFTD.^22,113,121^ Interestingly, PSP typically shows greater involvement of subcortical and brainstem structures that connect to mesocortical and isocortical areas via long-range pyramidal circuits,^122^ which may have contributed to patterns of cortical gradient loss we observed. Larger cohorts of these rare subgroups of bvFTD are needed to confirm these patterns of neurodegeneration.

### LIMITATIONS AND FUTURE DIRECTIONS

Our is study is the largest histologic investigation of neurodegeneration in bvFTD to date but was limited by the examination of only two neuronal markers and single sections from five cytoarchitectonic types within the frontal lobe. However, we validated our digital metrics of NeuN-ir and SMI32-ir by showing that the laminar and regional data produce distributions consistent with laminar cytoarchitecture and cortical gradients of the frontal lobe in HC. Due to the evolution of clinical consensus criteria and testing batteries, we chose to assess behavioral symptoms in a consistent manner via retrospective chart review and used harmonized neuropsychological testing data available in subsets of our total cohort. While we encountered some missing data due to sample availability or the poor integrity of staining or tissue, we importantly used LME models to account for missing data and showed sample sizes per analysis for transparency. It is possible that sampling of additional cytoarchitectonic types and neuronal markers may identify other novel distributions of neurodegeneration in cortical gradients present in other lobes of the brain that likely contribute to the clinical profile of bvFTD. Indeed, our recent comparative case report of a PiD patient and TDP-C patient found relatively distinct patterns of neurodegeneration that followed medial-to-lateral and anterior-to-posterior cortical gradients in frontal and temporal lobes.^19^ Patterns of neuron loss along additional cortical gradients were outside the scope of the current investigation, but new high-throughput methods may support these anatomical investigations in more regions and cases. For example, novel segmentation and machine learning approaches are under development and will be deployed in future studies to detect, classify, and quantify specific classes of neurons and protein inclusions based on morphology, size, and other neurochemical signatures.

In conclusion, we used the cortical gradient as a unique neuroanatomical framework to identify which cortical layers and cytoarchitectonic types are differentially susceptible to pyramidal neurodegeneration in bvFTD-tau relative to bvFTD-TDP. Long-range pathways may contribute to the location and greater extent of pyramidal neurodegeneration in bvFTD-tau, highlighting the anatomical heterogeneity in bvFTD and the possibility that pathologic changes to separate cortical layers and regions may converge downstream onto common networks that lead to clinically indistinguishable presentations such as bvFTD. While additional neurons, glia, and axonal connections systematically vary along cortical gradients in healthy brains, their patterns of change in FTLD-related disorders remain unexplored. Future studies may leverage the cortical gradient to bridge topographical patterns between histologic and neuroimaging studies, leading to new insights into neuronal vulnerability, resilience, pathologic spread, and neural correlates of cognitive decline.

## ACKNOWLEDGEMENTS

We extend our gratitude and remembrance to Dr. Murray Grossman and Dr. John Trojanowski whose legacies to FTD research are far reaching and deeply impactful. We greatly appreciate the technical assistance provided by Winifred Trotman, Emily Xie, Alejandra Bahena, John Robinson, and Theresa Schuck. We also thank the patients and families for their invaluable contributions and participation in the brain donation program that made this study possible.

## SUPPLEMENTARY MATERIAL

Supplementary material is available online.

## ETHICS APPROVAL

All procedures performed in studies involving human participants were in accordance with the ethical standards of University of Pennsylvania Internal Review Board and with the 1964 Helsinki declaration and its later amendments or comparable ethical standards. This article does not contain any studies with animals performed by any of the authors.

## INFORMED CONSENT

Informed consent was obtained from all individual participants included in the study.

## AUTHOR CONTRIBUTIONS

All authors contributed to the interpretation of the work, revisions of the manuscript, and approved the submitted manuscript. DTO and DJI contributed to the design, data acquisition, analyses, interpretation, drafting, and revision of the work. SXX, NC, KAQC, and SA contributed to data acquisition and analyses.

