## Supplementary Material for "Cytoarchitectonic gradients of laminar degeneration in behavioral variant frontotemporal dementia"

| Cytoarchitectonic type | Topographic details | Cytoarchitectonic features |
| --- | --- | --- |
| **Type 1 (BA33)** | Periallocortex found in the fundus of the supracollasal sulcus adjacent to allocortical indusium griseum[62,65] | • Agranular (i.e., layer IV absence)<br>• Few differentiable layers<br>• Internopyramidal (i.e., largest pyramidal neurons found in infragranular layers) |
| **Type 2 (BA24a-c)** | Mesocortex/proisocortex that is comprised of BA24a, 24b, 24c and arranged along the dorsal-ventral axis of the cingulate gyrus:<br>• 24a is most ventral and borders **type 1** in callosal sulcus<br>• 24b borders 24a and 24c in the cingulate gyral crown<br>• 24c is most dorsal and borders **type 3** in postgenual cingulate sulcus[62-64] | • Agranular (i.e., layer IV absence)<br>• Supragranular and infragranular layers are differentiable but sublayer borders are irregular<br>• Internopyramidal (i.e., largest pyramidal neurons found in infragranular layers) |
| **Type 3 (BA32)** | Mesocortex/proisocortex found in paracingulate gyri/sulci where it borders 24c in cingulate and **type 4** in orbitofrontal [62-64] | • Dysgranular (i.e., thin, irregular layer IV)<br>• Irregular sublayer differentiation within supragranular and infragranular layers<br>• Largest pyramidal neurons mostly found in infragranular layers and sparsely found in layer III |
| **Type 4 (BA14/11)** | Isocortex found in the gyrus rectus and medial orbital gyri[29,30,56,66,67] | • Granular/eulaminate (i.e., six well-differentiated layers, including thicker, continuous layer IV)<br>• Largest pyramidal neurons found equally between supragranular and infragranular layers<br>• Differentiable sublayers of supragranular and infragranular layers |
| **Type 5 (BA46)** | Isocortex found in lateral gyri including the middle frontal gyrus[29,30,57,68] | • Granular/eulaminate (i.e., six well-differentiated layers, including thicker, continuous layer IV)<br>• Externopyramidal (i.e., largest pyramidal neurons more often found in supragranular layers)<br>• Sharper differentiation of sublayers in supragranular and infragranular layers |

**Supplementary Table 1: Topographic and cytoarchitectonic characteristics of each cytoarchitectonic type that guided reliable sampling.** BA, Brodmann area.

**A**

**NeuN**

| Main Groups | Total Patients | Hemisphere | Cytoarchitectonic Types | | | | | *Total* |
|---|---|---|---|---|---|---|---|---|
| | | | 1 | 2 | 3 | 4 | 5 | |
| **HC** | 32 | Left | 13 | 16 | 16 | 14 | 12 | *71* |
| | | Right | 9 | 13 | 14 | 14 | 8 | *58* |
| | | Left+Right | 22 | 29 | 30 | 28 | 20 | *129* |
| **bvFTD-TDP** | 47 | Left | 22 | 32 | 29 | 33 | 29 | *145* |
| | | Right | 15 | 24 | 18 | 28 | 24 | *109* |
| | | Left+Right | 37 | 56 | 47 | 61 | 53 | *254* |
| **bvFTD-tau** | 27 | Left | 13 | 15 | 12 | 12 | 13 | *65* |
| | | Right | 9 | 13 | 12 | 15 | 15 | *64* |
| | | Left+Right | 22 | 28 | 24 | 27 | 28 | *129* |

**SMI32**

| Main Groups | Total Patients | Hemisphere | Cytoarchitectonic Types | | | | | *Total* |
|---|---|---|---|---|---|---|---|---|
| | | | 1 | 2 | 3 | 4 | 5 | |
| **HC** | 32 | Left | 13 | 16 | 16 | 14 | 12 | *71* |
| | | Right | 9 | 13 | 14 | 14 | 8 | *58* |
| | | Left+Right | 22 | 29 | 30 | 28 | 20 | *129* |
| **bvFTD-TDP** | 47 | Left | 21 | 30 | 29 | 34 | 32 | *146* |
| | | Right | 16 | 24 | 18 | 27 | 21 | *106* |
| | | Left+Right | 37 | 54 | 47 | 51 | 53 | *242* |
| **bvFTD-tau** | 27 | Left | 12 | 15 | 12 | 12 | 14 | *65* |
| | | Right | 11 | 14 | 12 | 17 | 12 | *66* |
| | | Left+Right | 23 | 29 | 24 | 29 | 26 | *131* |

**B**

**NeuN**

| Pathologic Subgroups | | Total Patients | Hemisphere | Cytoarchitectonic Types | | | | | *Total* |
|---|---|---|---|---|---|---|---|---|---|
| | | | | 1 | 2 | 3 | 4 | 5 | |
| **bvFTD-TDP** | ***TDP-A*** | 20 | Left+Right | 14 | 25 | 23 | 30 | 25 | 117 |
| | ***TDP-B*** | 15 | Left+Right | 12 | 16 | 14 | 18 | 15 | 75 |
| | ***TDP-C*** | 6 | Left+Right | 5 | 7 | 4 | 7 | 5 | 28 |
| | ***TDP-E*** | 6 | Left+Right | 6 | 8 | 6 | 6 | 8 | 34 |
| **bvFTD-tau** | **CBD** | 4 | Left+Right | 4 | 5 | 5 | 5 | 5 | 24 |
| | **PiD** | 12 | Left+Right | 8 | 12 | 10 | 12 | 12 | 54 |
| | **PSP** | 5 | Left+Right | 5 | 5 | 5 | 5 | 6 | 26 |
| | **UTau** | 6 | Left+Right | 5 | 6 | 4 | 5 | 5 | 25 |

**SMI32**

| Pathologic Subgroups | | Total Patients | Hemisphere | Cytoarchitectonic Types | | | | | *Total* |
|---|---|---|---|---|---|---|---|---|---|
| | | | | 1 | 2 | 3 | 4 | 5 | |
| **bvFTD-TDP** | ***TDP-A*** | 20 | Left+Right | 14 | 24 | 23 | 29 | 26 | 116 |
| | ***TDP-B*** | 15 | Left+Right | 13 | 16 | 14 | 18 | 14 | 75 |
| | ***TDP-C*** | 6 | Left+Right | 5 | 7 | 4 | 7 | 6 | 29 |
| | ***TDP-E*** | 6 | Left+Right | 5 | 7 | 6 | 7 | 7 | 32 |
| **bvFTD-tau** | **CBD** | 4 | Left+Right | 4 | 5 | 5 | 5 | 5 | 24 |
| | **PiD** | 12 | Left+Right | 9 | 12 | 9 | 12 | 10 | 52 |
| | **PSP** | 5 | Left+Right | 5 | 5 | 6 | 6 | 5 | 27 |
| | **UTau** | 6 | Left+Right | 5 | 7 | 4 | 6 | 6 | 28 |

| C | NeuN | | | | | | | | |
|---|---|---|---|---|---|---|---|---|---|
| | **Main Groups** | **Total Patients** | **Hemisphere** | **Cytoarchitectonic Types** | | | | | ***Total*** |
| | | | | **1** | **2** | **3** | **4** | **5** | |
| | **HC** | 32 | Bilateral | 0 | 0 | 0 | 0 | 0 | *0* |
| | **bvFTD-TDP** | 47 | Bilateral | 3 | 8 | 8 | 13 | 8 | *40* |
| | **bvFTD-tau** | 27 | Bilateral | 1 | 2 | 3 | 2 | 2 | *10* |

| SMI32 | | | | | | | | |
|---|---|---|---|---|---|---|---|---|
| **Main Groups** | **Total Patients** | **Hemisphere** | **Cytoarchitectonic Types** | | | | | ***Total*** |
| | | | **1** | **2** | **3** | **4** | **5** | |
| **HC** | 32 | Bilateral | 0 | 0 | 0 | 0 | 0 | *0* |
| **bvFTD-TDP** | 47 | Bilateral | 4 | 7 | 8 | 14 | 8 | *41* |
| **bvFTD-tau** | 27 | Bilateral | 3 | 3 | 2 | 3 | 0 | *11* |

**Supplementary Table 1: Frequency of cytoarchitectonic types used in analyses of NeuN-ir and SMI32-ir.**

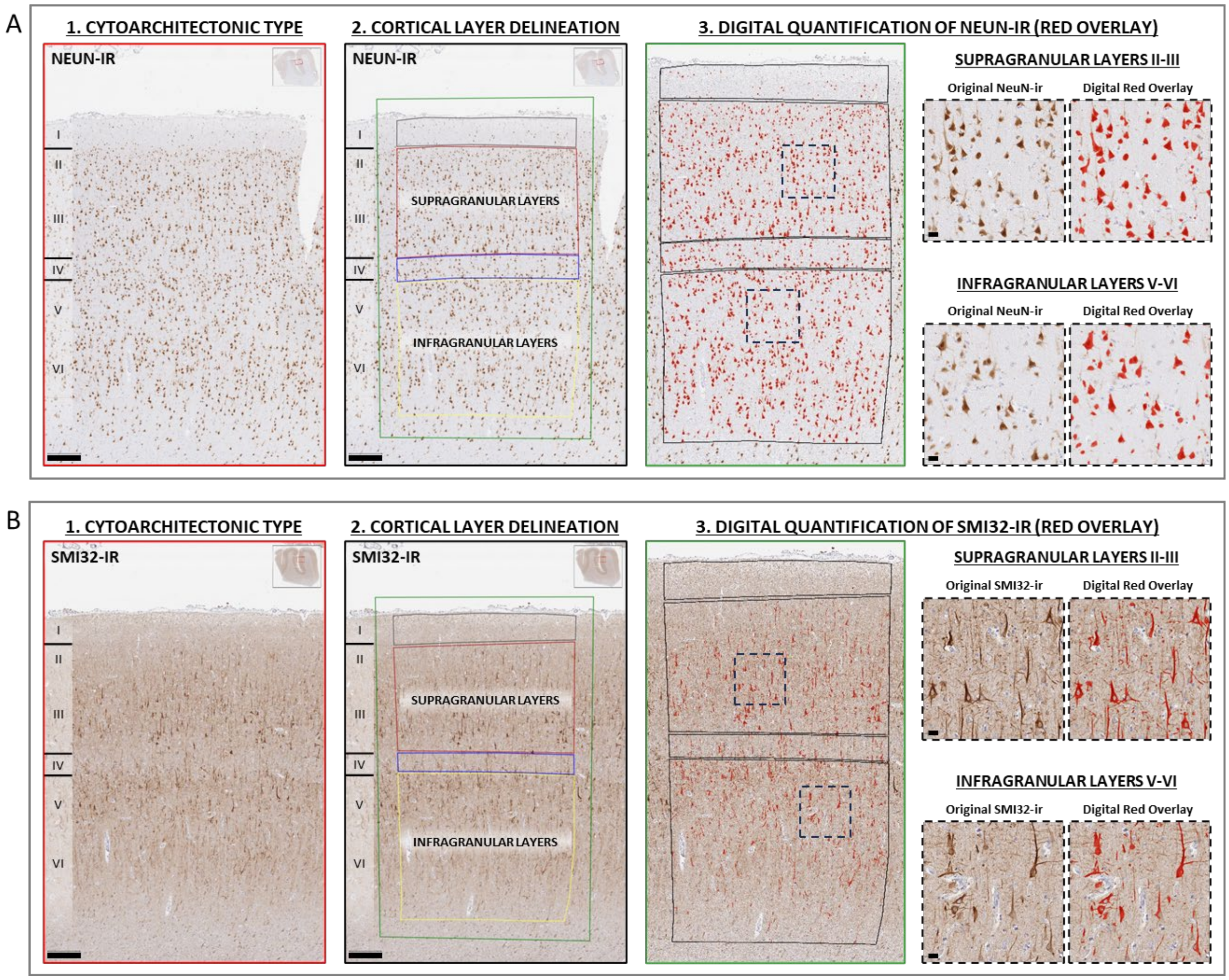


**Supplementary Figure 1: Digital quantitation of neuronal markers NeuN and SMI32 within delineated laminar subregions.**

We followed a strict protocol to identify cytoarchitectonic types of interest (**Supp. Table 1**) and then delineate cortical layers in each cytoarchitectonic type before digitally quantifying the % area occupied by NeuN-ir (**Supp. Fig. 1A**) and SMI32-ir (**Supp. Fig. 1B**) per section. Extensive literature demonstrates that SMI32 is a sensitive marker of human cytoarchitecture and cortical gradients due to its reliable labeling of non-phosphorylated neurofilament-rich pyramidal neurons.[27,37,55,68,69] Therefore, SMI32-ir was the most critical determinant of cortical layers and cytoarchitectonic types in the present study, but cytoarchitecture was assessed using all neuronal stained tissue. For example, in severely degenerated tissue with minimal SMI32-ir, densities of hematoxylin-positive cells and principles of topology/topography helped differentiate cytoarchitectonic types and cortical layers.

In brief, the first step involved identifying representative areas of laminar cytoarchitecture from the whole slide image, typically in flat areas where cortical layers were readily discernible (red border panels). In the second step, an experienced researcher (D.T.O.) used QuPath to manually delineate layer I, supragranular layers II-III, layers IV, and infragranular layers V-VI in each cytoarchitectonic type available per section (black border panels). Laminar subregions corresponded closely between semi-adjacent sections with NeuN-ir and SMI32-ir (**Supp. Fig. 1A compared to Supp. Fig. 1B**). The third and last step involved digital quantification of immunoreactivity that included detailed methods as follows:

First, we used an adaptive thresholding approach to digitally remove non-specific background. Whole-slide images were then exported as json files to overlay a grid of adjacent tiles (1024x1024 pixels each) restricted to pixels inside of annotated cortical layers only. The non-specific background is estimated via convolving a gaussian kernel of size 200x200 microns over the image, which is then subtracted from the image. A threshold is then selected for this normalized image through a maximum deviation thresholding algorithm. The thresholding algorithm uses the histogram of the normalized signal and an elbow method to select an optimal threshold for each ROI.

We quantified immunoreactive pixels (i.e., positive pixels [PosPix]) per area of interest using the calculation of percent area occupied (%AO) by immunoreactivity and visualized as red overlay in photomicrographs (green border panels and black dashed border panels). The %AO by NeuN-ir or SMI32-ir in cortical layers combined within cytoarchitectonic types or between subregions of cytoarchitectonic types was calculated as a weighted average (i.e., $\%AO^{WA}$) as follows:

$$\%AO^{WA} = \frac{\text{PosPix}(1) + \ldots + \text{PosPix}(n)}{\text{Area}(1) + \ldots + \text{Area}(n)} \text{ X } 100$$

In the current study, areas of interest refer to either distinct cortical layers or combined cortical layers within cytoarchitectonic types. For example, to measure the %AO across combined cortical layers I-VI within a given cytoarchitectonic type, we summed all positive pixels across cortical layers I-VI before dividing it by the summation of all pixels occupying cortical layers I-VI.

Similarly, we calculated the $\%AO^{WA}$ for supragranular and infragranular layers that were sampled more than once for a given cytoarchitectonic type (i.e., BA24a-c in cytoarchitectonic type 2, post-genual BA32 and subgenual BA32 in cytoarchitectonic type 3, and BA14-11 in cytoarchitectonic type 4). For example, we sampled layers of agranular mesocortex up to three times in the same section given that BA24 is composed of multiple spatially distinct subregions of the cingulate gyrus (i.e., 24a, 24b, 24c). Therefore, to calculate the $\%AO^{WA}$ in supragranular layers of agranular mesocortex, we divided the sum of the positive pixels across supragranular layers of 24a + 24b + 24c by the summed areas of supragranular layers in 24a + 24b +24c.

High magnification insets demonstrate the close correspondence between somatodendritic immunostaining of NeuN and SMI32 and the digitally measured area occupied of each neuronal marker (black dashed borders). All images were acquired from MFC sections of HC, representing the eulaminate-II isocortical type of cytoarchitecture. Low magnification scale bars = 250μm. High magnification scale bars = 20μm.

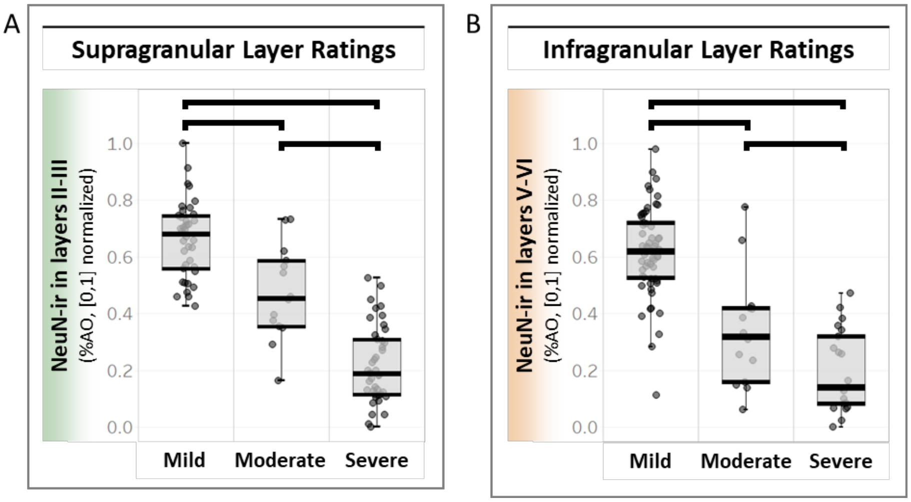


<u>Supplementary Figure 2:</u> **Validation of digital quantitation of NeuN-ir in cortical layers.**
To validate the digitally measured %AO of NeuN-ir, an experienced researcher (D.T.O.) using a 3-point rating scale to rank the level of NeuN immunostaining in supragranular layers and infragranular layers in all available MFC tissue across groups, blinded to group category. Mild, moderate, and severe ratings correspond to relatively high, intermediate, and low NeuN-ir in respective layers examined in both supragranular layers (**Supp. Fig. 2A**) and infragranular layers (**Supp. Fig. 2B**). LME models stratified by supragranular layers and infragranular layers found good agreement between the digitally measured NeuN-ir and ordinal ratings of NeuN-ir. We found digital measures of NeuN-ir were significantly different between ordinal ratings of NeuN-ir in both supragranular layers and ($F(2)=94.355$, $p<0.001$) and infragranular layers ($F(2)=57.305$, $p<0.001$). In post-hoc pairwise comparisons of ordinal ratings, we found a significant difference in digitally measured NeuN-ir between each ordinal rating in both supragranular layers and infragranular layers ($p<0.05$, Bonferroni corrected). These results suggest that digitally measured NeuN-ir accurately reflect the severity of overall neuron loss examined by cytoarchitectonic type and cortical layer.

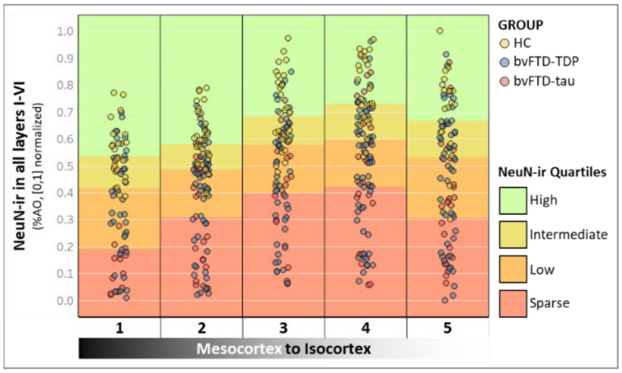


**Supplementary Figure 3: Stages of overall neurodegeneration severity assigned to each cytoarchitectonic type.** For each cytoarchitectonic type, we calculated high, intermediate, low, and sparse quartiles for NeuN-ir from all groups combined, reflecting stages of overall neurodegeneration (i.e., minimal, mild, moderate, severe). We staged neurodegeneration by cytoarchitectonic type to address the fact that neuron densities vary by regions of the cortical gradient in HC[37,70-72] and pathologic burden varies between regions in the FTLD spectrum.[18]

| **Upper Quartile of High NeuN-ir (Minimal Overall Neurodegeneration)** | | | | | | | |
|---|---|---|---|---|---|---|---|
| **Main Groups** | | **Cytoarchitectonic Types** | | | | | |
| | | **1** | **2** | **3** | **4** | **5** | ***Total*** |
| **HC** | | 11 | 17 | 17 | 17 | 13 | *75* |
| **bvFTD-TDP** | | 7 | 8 | 7 | 6 | 8 | *36* |
| **bvFTD-tau** | | 2 | 2 | 1 | 6 | 3 | *14* |

| **Upper Median Quartile of Intermediate NeuN-ir (Mild Overall Neurodegeneration)** | | | | | | | |
|---|---|---|---|---|---|---|---|
| **Main Groups** | | **Cytoarchitectonic Types** | | | | | |
| | | **1** | **2** | **3** | **4** | **5** | ***Total*** |
| **HC** | | 9 | 10 | 8 | 9 | 5 | *41* |
| **bvFTD-TDP** | | 6 | 12 | 11 | 15 | 14 | *58* |
| **bvFTD-tau** | | 4 | 5 | 5 | 3 | 5 | *22* |

| **Lower Median Quartile of Low NeuN-ir (Moderate Overall Neurodegeneration)** | | | | | | | |
|---|---|---|---|---|---|---|---|
| **Main Groups** | | **Cytoarchitectonic Types** | | | | | |
| | | **1** | **2** | **3** | **4** | **5** | ***Total*** |
| **HC** | | 2 | 2 | 5 | 2 | 1 | *12* |
| **bvFTD-TDP** | | 13 | 16 | 11 | 17 | 14 | *71* |
| **bvFTD-tau** | | 4 | 9 | 9 | 8 | 9 | *39* |

| **Lower Quartile of Sparse NeuN-ir (Severe Overall Neurodegeneration) *** | | | | | | | |
|---|---|---|---|---|---|---|---|
| **Main Groups** | | **Cytoarchitectonic Types** | | | | | |
| | | **1** | **2** | **3** | **4** | **5** | ***Total*** |
| **HC** | | 0 | 0 | 0 | 0 | 1 | *1* |
| **bvFTD-TDP** | | 9 | 16 | 16 | 19 | 13 | *73* |
| **bvFTD-tau** | | 11 | 11 | 9 | 9 | 10 | *50* |

**Supplementary Table 3: Frequency of cytoarchitectonic types assigned stages of total neurodegeneration severity.** *Due to lack of cytoarchitectonic types in the lower quartile for HC, we combined cytoarchitectonic types in lower and lower median quartiles for the analysis of SMI32-ir in cytoarchitectonic types with moderate-to-severe neurodegeneration (**Fig. 6B**).

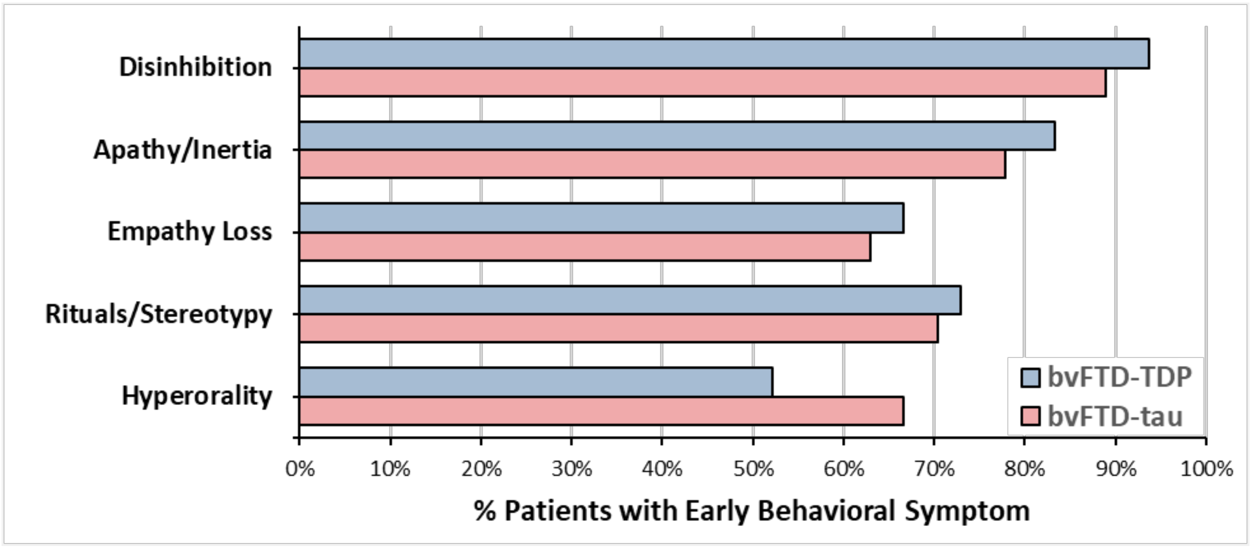


**Supplementary Figure 4: Similar prevalence of early core behavioral symptoms associated with bvFTD in bvFTD-TDP vs bvFTD-tau.** Presence and absence of behavioral symptoms within three years of symptom onset were evaluated by core diagnostic categories per bvFTD clinical criteria.[1] Behavioral symptoms were considered present if recorded at least once; symptoms were considered absent if not explicitly mentioned or could not be extrapolated from the records.

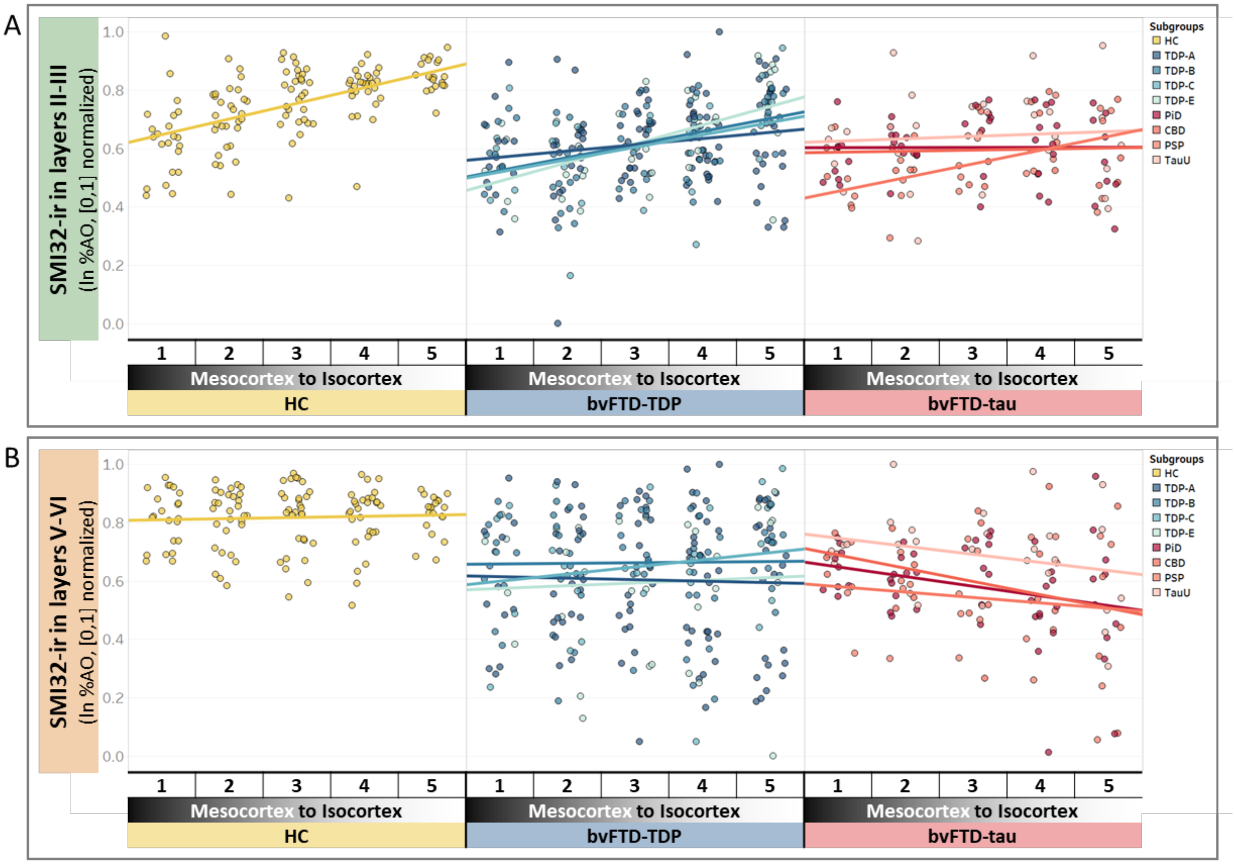


**<u>Supplementary Figure 5:</u> Distinct relationships between the cortical gradient and laminar pyramidal neurodegeneration in pathologic subgroups of bvFTD-TDP and bvFTD-tau.**

While statistical analyses were not performed in pathologic subgroups due to sample sizes, we evaluated the consistency of relationships between the cortical gradient and laminar SMI32-ir within main groups of bvFTD-TDP and bvFTD-tau as follows:

*Supragranular SMI32-ir findings*

Similar to main results found in bvFTD-TDP (**Fig. 4**), we find all pathologic subgroups of bvFTD-TDP maintain positive relationships between the cortical gradient and supragranular SMI32-ir (**Supp. Fig. 5A**). Similar to main results found in bvFTD-tau (**Fig. 4**), we find most pathologic subgroups of bvFTD-tau (i.e., PiD, CBD, and TauU) share no relationships between the cortical gradient and supragranular SMI32-ir (**Supp. Fig. 5A**). However, PSP showed a positive relationship between the cortical gradient and supragranular SMI32-ir, suggesting that PSP may have a predilection for supragranular pyramidal neurodegeneration in mesocortices compared to isocortices.

*Infragranular SMI32-ir findings*

Similar to main results found in bvFTD-TDP (**Fig. 4**), we find all pathologic subgroups of bvFTD-TDP show no relationships between the cortical gradient and infragranular SMI32-ir (**Supp. Fig. 5B**). Similar to main results found in bvFTD-tau (**Fig. 4**), we find all pathologic subgroups of bvFTD-tau share negative relationships between the cortical gradient and infragranular SMI32-ir (**Supp. Fig. 5B**).

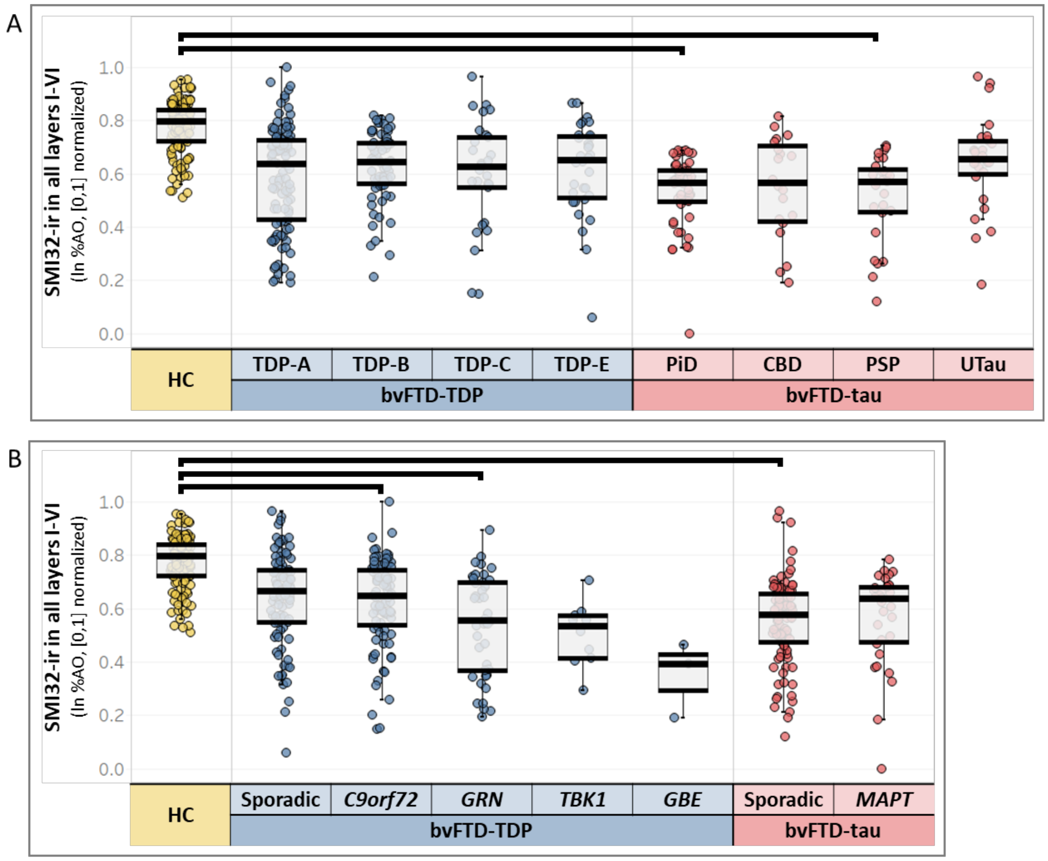


**<u>Supplementary Figure 6.</u> Patterns of neurodegeneration in bvFTD subgroups and HC.**
We find a significant difference in SMI32-ir between pathologic subgroups of bvFTD and HC (F[8,90.3]=4.039, p<0.001) (**Supp. Fig. 6A**). Posthoc pairwise comparisons found similar levels of SMI32-ir between pathologic subgroups of bvFTD, but PiD (β=0.168, SE=0.045, p=0.012) and PSP (β=0.242, SE=0.065, p=0.012) each had lower SMI32-ir compared to HC. We find a significant difference in SMI32-ir between mutation carrier subgroups of bvFTD, sporadic subgroups of bvFTD, and HC (F[7]= 4.291, p<0.001) (**Supp. Fig. 6B**). Posthoc pair-wise comparisons found *C9orf72* (β=0.13, SE=0.04, p=0.045), *GRN* (β=0.174, SE=0.054, p=0.044), and sporadic bvFTD-tau (β=0.169, SE=0.038, p<0.001) each had lower SMI32-ir compared to HC. Bars signify p<0.05 with Bonferroni correction for multiple comparisons.
